# Bendless-mediated K63 ubiquitination modulates cellular signalling to regulate *Drosophila* hematopoiesis

**DOI:** 10.1101/2025.09.05.674504

**Authors:** Asfa Kamal, Yash Sheregare, Chaitanya Patkar, Mrunmayi Markam, Rujul Deolikar, Nawaneetan Sriraman, Aditya Seth, Manish Jaiswal, Rohan Jayant Khadilkar

**Affiliations:** Stem Cell and Tissue Homeostasis laboratory, Advanced Centre for Treatment, Research and Education in Cancer (ACTREC), Tata Memorial Centre, Kharghar, Navi Mumbai, Maharashtra 410210, India; Homi Bhabha National Institute, Training School Complex, Anushakti Nagar, Mumbai, 400085, India; Tata Institute of Fundamental Research Hyderabad, Hyderabad, India

**Keywords:** K63 ubiquitination, Cellular Signalling, Homeostasis, Hematopoiesis, *Drosophila*

## Abstract

Ubiquitination is a reversible modification whose traditional role has been associated with K48-linked poly-ubiquitination involved in proteasomal degradation. However, the role of K63-linked poly-ubiquitination has been explored in various cellular processes like DNA repair, endocytosis, innate immune response, kinase activation, etc. Since K63-linked poly-ubiquitination can regulate the stability, localization, and activity of its target molecules, its regulation and function in various developmental and disease contexts are being explored. Here, we investigate how K63 ubiquitination regulates *Drosophila* blood cell homeostasis. Ubc13 (UBE2N), an E2 conjugating enzyme, is highly expressed in Acute Myeloid Leukemia (AML), wherein it controls innate immune signalling for the survival of AML cells; however, its role during developmental hematopoiesis is less explored. In *Drosophila*, Bendless is the functional homolog of Ubc13, whose role in stem cell regulation and particularly hematopoiesis is unknown. Our results indicate that spatial genetic perturbation of Bendless and its associated molecules, namely Uev1a, Effete, and E3 ligase - Traf6, that mediate K63 ubiquitination are critical for maintaining hematopoietic progenitors and regulating their differentiation. We show that Bendless-mediated K63 ubiquitination controls the Wingless signalling pathway by regulating Dishevelled in the larval lymph gland (LG) thereby modulating hematopoietic progenitor maintenance and differentiation. Furthermore, an excess of K63 ubiquitination activates the JNK pathway in the LG, resulting in lamellocyte production. Genetic epistasis analysis shows that activation of the canonical Wingless pathway in the background of Bendless depletion or inactivation of the JNK pathway in Bendless over-expression conditions can restore physiological hematopoiesis. Our findings indicate that regulators of K63 ubiquitination, like Bendless, could act as molecular inter-nodes that are capable of signalling cross-regulation, especially where intricate signalling networks are involved. Our study provides important mechanistic insights into the signalling mechanisms regulated by K63 ubiquitination during stem cell homeostasis.

## Introduction

Ubiquitin (Ub) is a small, highly conserved 76-amino acid protein found in all eukaryotic cells. It is covalently attached to substrate proteins through a sequential enzymatic cascade involving three types of enzymes: Ub-activating enzymes (E1), Ub-conjugating enzymes (E2), and Ub ligases (E3). This cascade begins with the formation of a thioester bond between Ub and the E1 enzyme, followed by transfer of Ub to a cysteine residue on the E2 enzyme. With the assistance of an E3 ligase, Ub is finally conjugated to the ε-amino group of lysine residues on target proteins (1). Ubiquitylation is a dynamic and reversible post-translational modification, as deubiquitylating enzymes (DUBs) can remove Ub from substrates or process Ub precursors (2). Proteins can undergo monoubiquitylation, multi-monoubiquitylation, or polyubiquitylation, wherein Ub molecules form chains via any of their seven lysine residues—K6, K11, K27, K29, K33, K48, and K63 (3). Traditionally, polyubiquitination was associated with proteasomal degradation, particularly through K48-linked chains (1) however K63-linked ubiquitination, the second most common form of ubiquitination in yeast and mammalian cells, is now well recognized for its non-proteolytic roles, including regulation of endocytosis, kinase activation, DNA repair, transcription, and innate immune responses (3,4). This form of ubiquitination is particularly important in modulating signaling pathways like NF-κB and JNK, thereby influencing processes such as immune response, inflammation, and cell fate decisions (5,6,7).

In *Drosophila*, *bendless* (ben) encodes an E2 ubiquitin-conjugating enzyme homologous to mammalian Ubc13, and it plays a crucial role in catalyzing K63-linked polyubiquitination (8). Ben functions with Uev1a to assemble non-degradative ubiquitin chains that regulate various cellular pathways, particularly in escape response (9), axon guidance (8), synaptic growth and maturation (10), long-term memory (11), innate immunity (5) and genomic integrity (12). Although the role of Bendless in neural development, immune signaling, and genomic stability is well-documented, its function in physiological hematopoiesis remains unexplored. This represents a significant gap in our understanding, especially given that its mammalian ortholog Ubc13 is essential for hematopoietic stem cell (HSC) function, as shown by studies in mice where Ubc13 deletion leads to loss of immune cells, thymic and bone marrow atrophy, and early lethality (13). Apart from the role of Ubc13 in developmental hematopoiesis, recent work has shown that AML hematopoietic stem and progenitor cells (HSPCs) have dysregulated innate immune signalling that relies on UBE2N (Ubc13) for leukemic cell function and inhibition of this enzyme selectively impairs leukemic cells (14–15). These findings highlight an important role of Ubc13 mediated K63 ubiquitination in regulating both physiological and malignant hematopoiesis. However, K63 ubiquitination mediated regulation of signalling mechanisms that control hematopoietic stem cell maintenance and fate decisions remain less explored.

*Drosophila* hematopoiesis occurs in two temporally distinct phases: an initial wave during embryogenesis and a second, definitive wave during larval development where the lymph gland (LG), serves as the primary hematopoietic site (16–17). The larval lymph gland is organized into functionally distinct developmental zones. The Posterior Signaling Center (PSC) acts as a hematopoietic niche, producing cues that not only support the maintenance of prohemocytes but also prime them towards differentiation (18–20). The Medullary Zone (MZ) contains undifferentiated progenitor cells capable of giving rise to the differentiated blood cell types namely — plasmatocytes, crystal cells, and lamellocytes that reside in the Cortical Zone (CZ) (17). Recent advances in single-cell transcriptomics have revealed substantial heterogeneity among lymph gland hemocytes, identifying previously unrecognized subpopulations such as adipohemocytes, core, distal and intermediate progenitors, and early lamellocytes, underscoring the complexity and dynamic organization of this hematopoietic tissue (21). These studies suggest that the lymph gland harbors a diverse set of hemocyte subsets governed by tightly regulated signaling networks (21–22).

The cell fate decisions of prohemocytes residing in the MZ are precisely controlled by both extrinsic and intrinsic signals (18, 20, 23–28, 29–30). Several evolutionarily conserved signaling pathways, including JAK/STAT, Hedgehog (Hh), Wingless (Wg), and Decapentaplegic (Dpp), have been implicated in the regulation of progenitor maintenance and differentiation within the LG (20, 23–24, 31–32, 33–34). Since the hematopoietic progenitors are tightly regulated by an intricate signalling network that consists of nodes and internodes that are capable of cross-talk and cross-regulation, understanding the on-off switches that modulate various signals becomes important. K63 ubiquitination could be one such context-dependent switch which needs further exploration.

Here, in this study we sought out to understand the role of K63 ubiquitination in the regulation of developmental hematopoiesis using Bendless as a tool. We explore the role of Bendless mediated K63 ubiquitination in regulating *Drosophila* larval LG hematopoiesis. *bendless* whole animal mutants show abrogated blood cell homeostasis and genetic perturbation of Bendless in the hematopoietic progenitors affects the progenitor population leading to increased blood cell differentiation. Our results show that Bendless functions with Uev1a, Effete and the E3 ligase, Traf6, in regulating hematopoiesis. We show that a critical balance of K63 ubiquitination is important for orchestrating hematopoiesis. We show that Bendless regulates the Wingless pathway to control progenitor maintenance and their subsequent differentiation, whereas it regulates lamellocyte differentiation via the JNK pathway. Taken together, we demonstrate that Bendless-mediated K63 ubiquitination levels are critical for maintaining signalling output in the blood cell progenitors in *Drosophila,* and a failure of maintenance of optimal levels of K63 ubiquitination skews differentiation trajectories. These results establish a novel role for Ben in regulating steady-state blood cell development and lay the foundation for further mechanistic studies into ubiquitin-mediated control of hematopoiesis in metazoans.

## Results

### Bendless is expressed in the Lymph Gland and Is Essential for Blood Cell Homeostasis in *Drosophila*

In *Drosophila*, *bendless* (ben) encodes an E2 ubiquitin-conjugating enzyme homologous to mammalian Ubc13 that catalyzes K63-linked polyubiquitination (8). Ben functions with Uev1a to assemble K63 ubiquitin chains that regulate various developmental processes described earlier (5, 8–11). Now, UBE2N (Ubc13) has been previously reported to be overexpressed in acute myeloid leukemias (AML) and is required for the survival of AML cells (35). The role of the *Drosophila* counterpart, Bendless in regulating various developmental processes has been explored, particularly in neuro-development; however, its role in regulating stem cell homeostasis remains to be understood. Based on the prior reports indicating an important role for its mammalian counterpart in leukemia, we set out to understand if Bendless mediated K63 ubiquitination controls developmental hematopoiesis in *Drosophila* (Fig. 1A). We first investigated whether Ub-K63 signals are observed in the larval LG and its hematopoietic progenitors. We detected the K63 linkages using an Ub-K63 antibody, and we found that the primary LG lobe, including the tepIV positive hematopoietic progenitors, has the presence of K63 chains (Fig. 1B-F). Bendless expression analysis using the antibody shows that Bendless is expressed in the LG hematopoietic progenitors (Fig. 1G-J). Single-cell RNA sequencing data mined from the fly hemocyte atlas shows that Bendless is expressed in almost all LG subsets, including the blood progenitor population (Fig. 1K). We also validated the specificity of the Bendless antibody and the knockdown and over-expression constructs of Bendless by using whole LG-specific *e33cGal4,* and our validation results indicate a reduction in Ben levels upon *e33cGal4-*mediated depletion of Ben, whereas an increase upon Ben over-expression (Fig. S2A-F).

**Figure 1:**
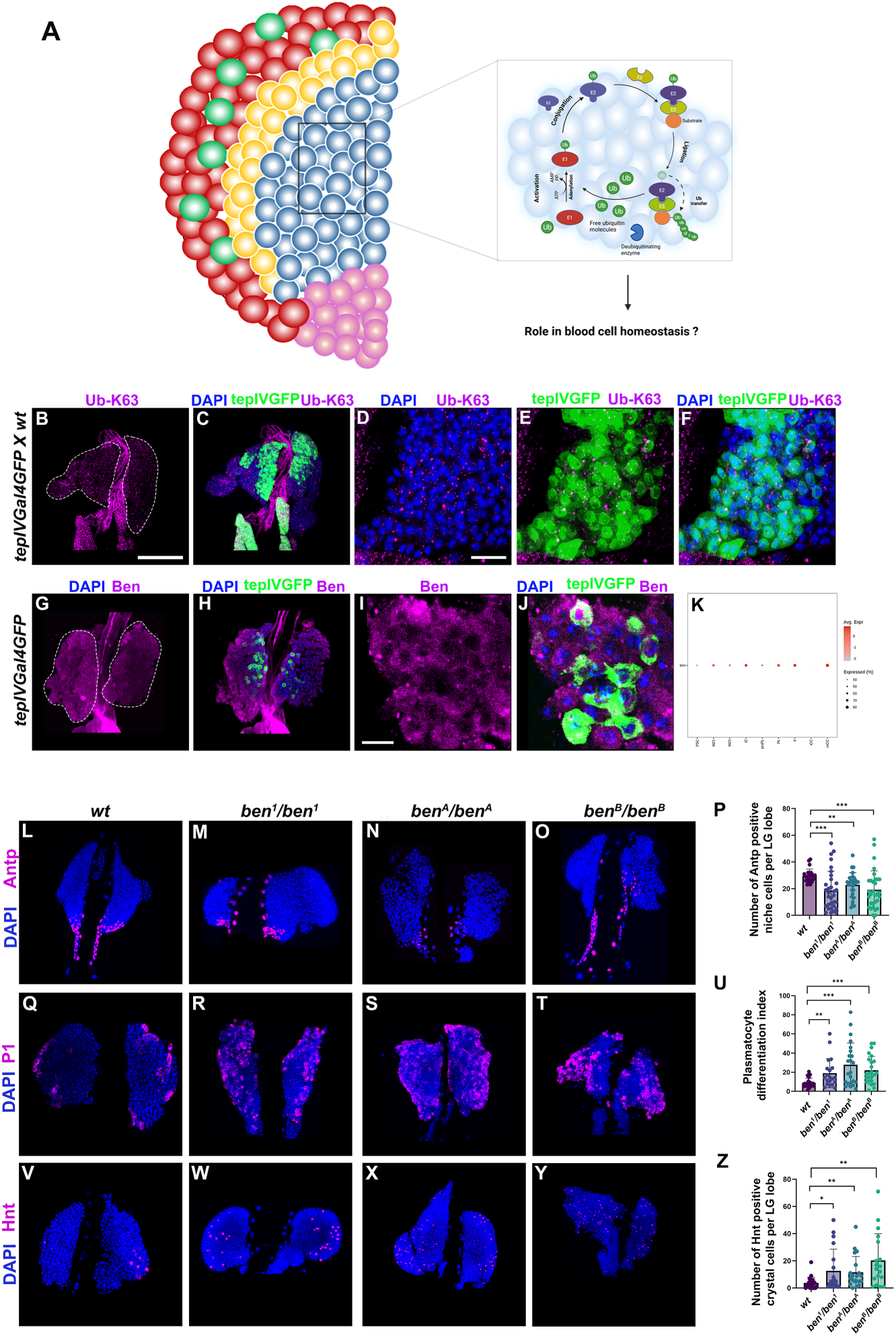
Bendless is expressed in the *Drosophila* larval lymph gland (LG) and regulates LG hematopoiesis. Schematic posing the question of how K63 ubiquitination could regulate blood cell homeostasis in the *Drosophila* LG (A). K63 ubiquitination pattern detected using anti-Ub-K63 antibody (magenta) in the entire primary LG lobe (B-C) and *tepIV-GFP* positive progenitors (Green) (D-F). Bendless expression (magenta) in the whole primary LG lobe (G-H) and *tepIV-GFP* positive progenitors (Green) (I-J). Plot from scRNA seq data from the Fly hemocyte atlas showing Bendless expression in various subsets of the LG (K). Posterior Signalling Centre/Niche cells marked with Antennapedia (Magenta) in LGs from *ben* mutant alleles (*ben^1^, ben^A^, ben^B^*) as compared to wild type control (L-O) with PSC cell numbers quantitated and represented graphically (P). Plasmatocyte differentiation marked with P1 (Magenta) in *ben* mutant alleles (*ben^1^, ben^A^, ben^B^*) as compared to wild type control (Q-T) with plasmatocyte differentiation index quantified and represented graphically (U). Crystal cell differentiation marked with Hindsight/Hnt (Magenta) in *ben* mutant alleles (*ben^1^, ben^A^, ben^B^*) as compared to wild type control (V-Y) with the number of crystal cells quantified and represented graphically (Z). Nuclei are marked with DAPI (Blue). GFP (Green) is driven by *tepIV-Gal4*. Scale Bar: 50µm (B-C, G-H, L-Y), 30µm (D-F, I-J). Statistical analysis was performed using Student’s t-test with Welch’s correction. For P, U, and Z, each data point represents data from a single LG lobe. P-values denoted are as follows: *** (P<0.001), ** (P<0.01), and * (P<0.1). Schematic: Created in BioRender. Khadilkar, R. (2025) https://BioRender.com/egilweh

Since Ben is expressed in all cellular subsets of the LG, we first wanted to examine the LG phenotypes of the *ben* mutant alleles. To do this, we analyzed the LG hematopoietic phenotypes in three independent EMS-generated amorphic alleles of *ben* (*ben^1^/ben^1^, ben^A^/ben^A^* and *ben^B^/ ben^B^*) in homozygous conditions. We found that the number of Antennapedia-positive Posterior Signaling Centre (PSC) niche cells were significantly reduced in all three mutants (*ben^1^/ben^1^, ben^A^/ben^A^, ben^B^/ ben^B^*) compared to the wild type (Fig. 1L-P). Interestingly, LGs of all the mutant alleles showed a significant increase in P1 (NimRodC1)-positive plasmatocyte population (Fig. 1Q-U) and Hindsight-positive crystal cells compared to wild type (Fig. 1V-Z). Along with these, we also observed lamellocyte-positive lymph glands in all three *ben* mutant alleles (Fig. S1A-I), even in the absence of parasitic wasp infestation. Since these are whole animal mutants where multiple tissues expressing Bendless could be affected, along with deregulation of systemic signalling, the LG may respond to such systemic cues by switching on its blood cell differentiation program.

### Hematopoietic progenitor-specific depletion of Bendless results in aberrant blood cell differentiation

Since the *ben* mutant LGs showed increased blood cell differentiation, we wanted to investigate the effect of Bendless depletion in LG hematopoietic progenitors. We used two different Gal4 drivers, namely *tepIV-Gal4* for core progenitors and *domeless-Gal4* for distal progenitors. Both *tepIV* (Fig. 2A-F) or *domeless-Gal4* mediated (Fig. 2G-L) depletion of Bendless led to an increase in P1 (NimRodC1 positive) plasmatocyte differentiation and Hindsight (Hnt) positive crystal cell differentiation. We also quantified the tepIV-GFP positive progenitor population upon Bendless depletion using *tepIV-Gal4* and found that the tepIV-GFP-positive progenitor population was reduced upon Bendless depletion as compared to the wild type control (Fig. S3A, B, D, E, and G).

**Figure 2:**
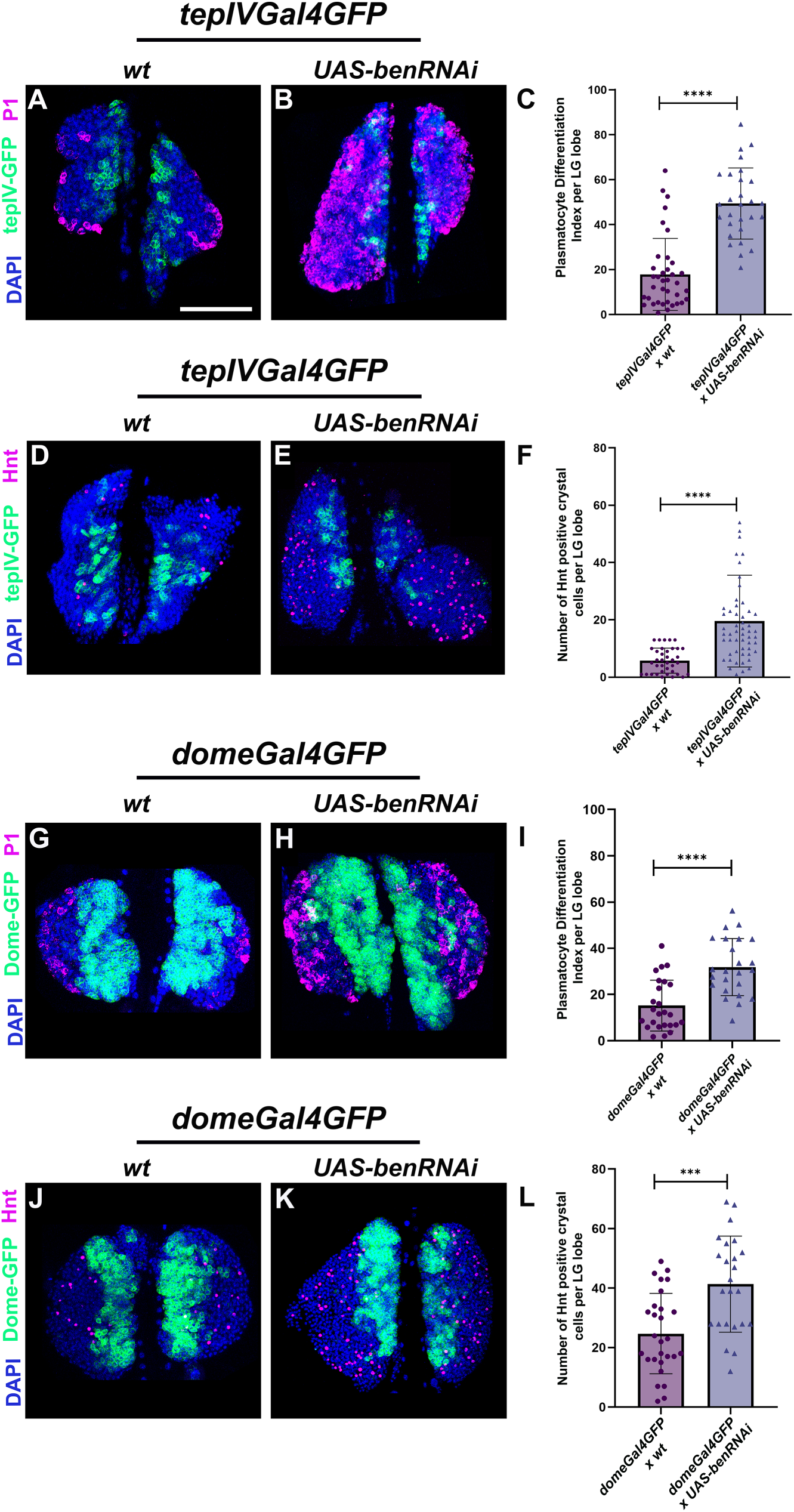
Modulation of Bendless levels in the hematopoietic progenitors alters blood cell homeostasis in the LG. Plasmatocyte differentiation marked by P1 (magenta) or crystal cell differentiation marked by Hindsight (Hnt) (magenta) upon expression of *UAS-benRNAi* in the core progenitors (using *tepIVGal4,* A-B, D-E) or distal progenitors (using *domeGal4,* G-H, J-K) as compared to respective *wildtype* control (A, D, G, and J). Graphical representation of plasmatocyte differentiation index or number of crystal cells upon *tepIVGal4* (C, F) or *domeGal4* (I, L) mediated Bendless knockdown as compared to *wildtype* control. GFP expression (green) is driven by *tepIVGal4* or *domeGal4*. Nuclei are stained with DAPI (Blue). Scale Bar: 50µm (A-B, D-E, G-H, J-K). Statistical analysis was performed using Student’s t-test with Welch’s correction. Each data point represents data from a single LG lobe. P-values denoted are as follows: **** (P<0.0001) and *** (P<0.001).

### Genetic perturbation of molecular regulators of K63 ubiquitination in the hematopoietic progenitors results in loss of blood cell homeostasis

Bendless depletion phenotypes prompted us to investigate whether perturbation of additional molecules participating in K63 ubiquitination show similar effects in the LG. To test this, we analyzed the role of Uev1a, a known partner of Ben in K63 ubiquitin chain assembly (36–37). Previous literature showed that Uev1a forms a stable heterodimer with Bendless (Ben), the *Drosophila* ortholog of Ubc13, to mediate K63-linked polyubiquitination (38). Disruption of this interaction through mutations in *dUev1a* impairs both the DNA damage response and neurological functions, underscoring the critical functional partnership between Uev1a and Ben (37). Knockdown of *uev1a* in tepIV positive progenitors resulted in increased blood cell differentiation phenotypes comparable to those observed in *ben* knockdown, including increased plasmatocyte and crystal cell differentiation compared to wild type (Fig. 3A-B, E-F, I-J). Effete (UbcD1) is another molecule, a highly conserved Class I E2 ubiquitin-conjugating enzyme in *Drosophila*, consisting solely of a UBC (ubiquitin-conjugating) catalytic domain essential for ubiquitin-mediated protein modification involved in various cellular processes, including eye development, germline stem cell maintenance, and apoptosis regulation (39–43). Interestingly, Effete depletion in tepIV positive progenitors led to a significant increase in the differentiation of both plasmatocytes and crystal cells as compared to wild type (Fig. 3A, C, E, G, I, and J). Furthermore, for the ubiquitination process to occur, an E3 ligase is required to confer substrate specificity and facilitate ubiquitin transfer. Previous reports suggest that TRAF6, a well-characterized E3 ubiquitin ligase, interacts with E2 enzymes to mediate K63-linked polyubiquitination (44). In *Drosophila*, the E2 enzyme Bendless (Ben) functions in the TNF-JNK pathway by interacting with dTRAF2, the fly ortholog of TRAF6, to regulate JNK activation, cell death, and tumor progression (7). To examine if TRAF6 depletion also phenocopies the phenotypes of Ben, Uev1a, and Effete depletion, we depleted TRAF6 in tepIV-positive progenitors. Knockdown of TRAF6 indeed resulted in a phenocopy effect wherein both plasmatocyte and crystal cell differentiation were increased in the LG lobe (Fig. 3A, D, E, H, I, and J). These data suggest that these molecules function along with Bendless in regulating blood cell homeostasis in the LG. Along with localized perturbation, we also studied the LG phenotype of whole animal mutants of *dtraf6.* We looked at the LG phenotypes of the *traf6^ex1^* amorphic allele, and our observations indicate that the PSC niche size decreases with a concomitant increase in both plasmatocyte and crystal cell differentiation, behaving very similarly to the *ben* mutant alleles (Fig. S4A-I).

**Figure 3:**
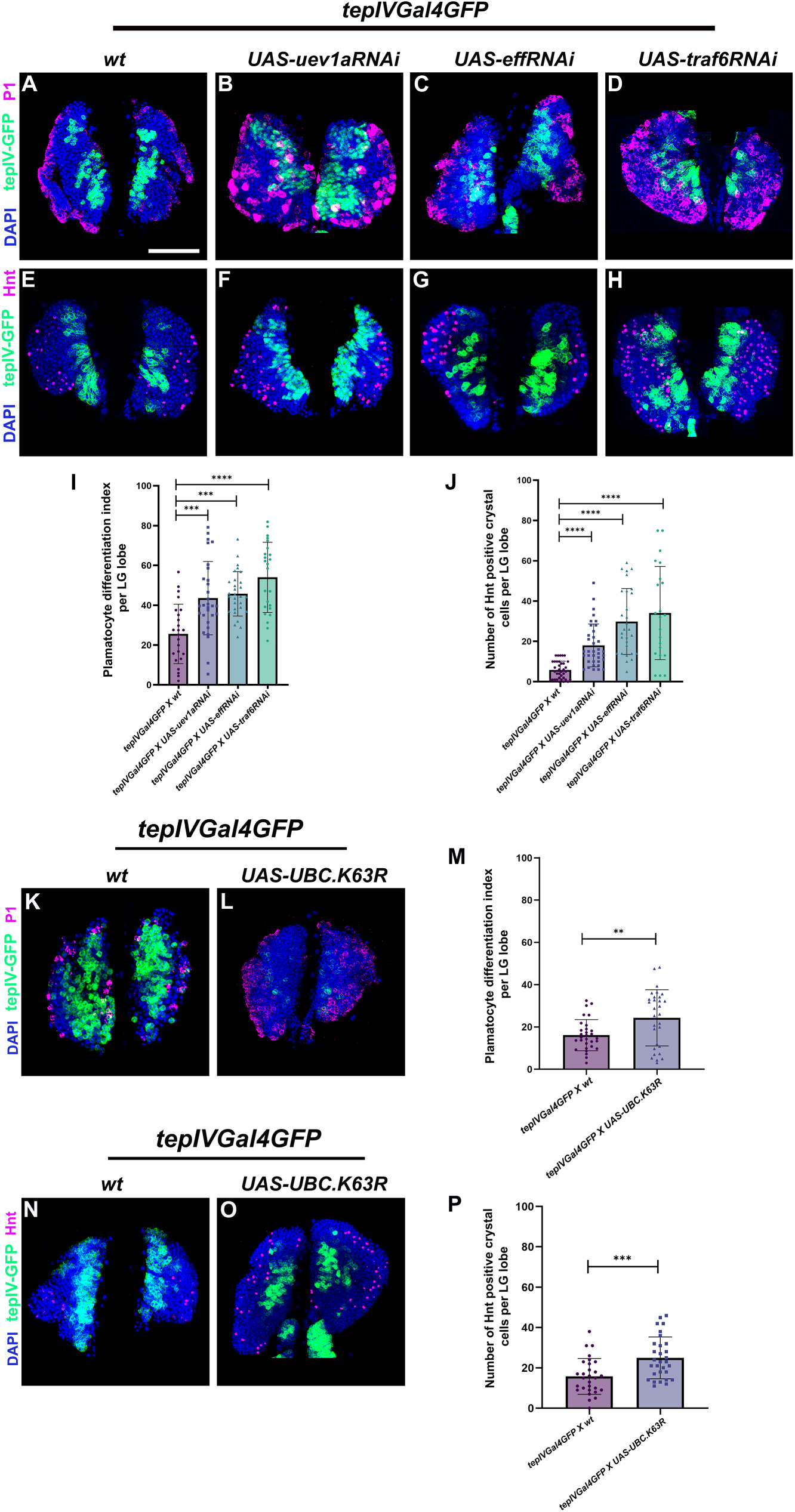
Genetic perturbation of molecules regulating K63 ubiquitination in core progenitors results in increased blood cell differentiation. Plasmatocyte differentiation marked by P1 (magenta) or crystal cell differentiation marked by Hnt (magenta) upon *tepIVGal4* mediated knockdown of *UAS-uev1aRNAi* (B, F), *UAS-effRNAi* (C, G) or *UAS-traf6RNAi* (D, H) or upon expression of a non-functional ubiquitin mutant, Ubc.K63R (L, O) as compared to their respective wildtype control (A, E, K, N). Graphical representation of Plasmatocyte differentiation index and total number of Hnt positive crystal cells per LG lobe upon expression of *UAS-uev1aRNA*i, *UAS-effRNAi*, or *UAS-traf6RNAi* or a non-functional ubiquitin mutant in the tepIV positive core progenitor population, compared to the wildtype (Fig. 3I, J, M, P). UASGFP or UAS-mCD8 expression (green) is driven by *tepIVGal4*. Nuclei are stained with DAPI (Blue). Scale Bar: 50µm (A-H, K-L, N-O). Statistical analysis was performed using Student’s t-test with Welch’s correction. Each data point represents data from a single LG lobe. P-values denoted are as follows: **** (P<0.0001), *** (P<0.001), and ** (P<0.01).

In addition to the above, we have also employed the *UAS-UBC.K63R* transgene to modulate K63 ubiquitination by expressing a ubiquitin variant in which lysine 63 is replaced by arginine (K63R) under UAS control, which leads to a defect in the formation of K63 ubiquitin linkages. We also validated this transgene using a K63-specific ubiquitin antibody and observed reduced signal (Fig. S5F-J) upon expression of the non-functional variant in tepIV positive progenitors as compared to the wild-type (Fig. S5A-E), confirming impaired K63-linked ubiquitination. Importantly, abrogation of K63 ubiquitination in the tepIV positive progenitors phenocopies loss of Ben phenotypes, leading to increased differentiation of both plasmatocyte (Fig. 3K-M) and crystal cell differentiation (Fig. 3N-P) along with a decrease in tepIV positive progenitor population (Fig. S3C, F, G) as compared to the wild type (Fig. S3A, D, G). Taken together, the phenotypic similarities caused by disruption of these components suggest that a conserved K63 ubiquitination module supports blood cell homeostasis, emphasizing the significance of ubiquitin-mediated signaling in hematopoietic control and providing a foundation for further mechanistic exploration of this pathway.

### Bendless-mediated K63 ubiquitination regulates Dishevelled, a component of the *Drosophila* Wingless pathway in the LG

Wingless (Wg)/Wnt signaling plays a vital role in maintaining prohemocytes in the *Drosophila* lymph gland by preventing their aberrant differentiation (24). Recently, it was also shown that Wnt6/EGFR pathways simultaneously control progenitor growth, proliferation, and their differentiation in the LG (45). Upon Wnt ligand binding to Frizzled and Lrp5/6 receptors, the cytoplasmic protein Dishevelled (Dvl/Dsh) is recruited to the receptor complex, where it facilitates the assembly of a signaling platform that inhibits the Axin-APC-GSK3β-CK1 complex. This inhibition stabilizes β-catenin, allowing its nuclear translocation and activation of Tcf/Lef-dependent transcription. Dsh thus serves as a key upstream regulator of canonical Wnt signaling by blocking β-catenin degradation and promoting downstream gene expression (46). Structurally, Dvl contains three conserved domains essential for its diverse signaling roles: the N-terminal DIX domain, which mediates homo- and heterodimerization; the central PDZ domain, which facilitates binding to Fz; and the C-terminal DEP domain, which engages in protein and lipid interactions (47–52).

Recent studies have highlighted the importance of ubiquitin modifications, particularly non-proteolytic K63-linked polyubiquitylation, in fine-tuning Wnt/β-catenin signaling (53). Ubiquitin chains, being inducible and reversible, serve as versatile signaling cues recognized by specific ubiquitin-binding domains (54). Notably, a general role for K63-linked ubiquitin chain formation in regulating Wnt signaling during hematopoiesis has been demonstrated using conditional Ubc13 knockout mice, underscoring the critical involvement of this modification in developmental signaling pathways (13). Now, there are multiple reports that indicate that Dishevelled gets K63 ubiquitinated, thereby regulating the outcome of the Wnt pathway. In the context of human cylindromatosis, it was shown that loss of CYLD, a deubiquitinase, leads to enhanced K63 ubiquitination of the DIX domain of Dvl, leading to active Wnt signalling (53) whereas another report suggests that Dvl2 forms condensates leading to phase separation upon ubiquitination by an E3 ligase named WWP2, resulting in active Wnt signalling (55).In contrast, another report suggested the opposite, where genetic or chemical suppression of Usp14, a deubiquitinase that acts on Dvl, leads to increased polyubiquitination of Dvl, resulting in impaired Wnt signalling (56).

Consistent with these findings, we hypothesized that Bendless regulates the differentiation of hematopoietic progenitors in the *Drosophila* lymph gland through the Wingless (Wg) signaling pathway, specifically via Dishevelled (Dsh), a key intracellular effector of canonical Wg signaling. To explore the regulation of this pathway, we first examined the co-localization of Dsh and K63-linked ubiquitin in tepIV-GFP positive hematopoietic progenitors and found that there is a significant co-localization between Ub-K63 and Dsh in the hematopoietic progenitors, indicating that Dsh may be getting K63 ubiquitinated in the LG (Fig. 4A-E’). Further supporting this, we observed reduced Dsh expression upon Bendless depletion in the tepIV-positive progenitors as compared to the control (Fig. 4F-I’’), aligning with previous reports that Ubc13-mediated K63-linked ubiquitination stabilizes Dsh. Additionally, expressing a non-functional Ubiquitin mutant that fails to form K63 linkages in the tepIV positive progenitors, too, led to a marked reduction in Dsh levels (Fig. 4J-M’’). To support our observations, we went ahead and depleted Dishevelled from the tepIV-GFP progenitors using *tepIV-Gal4,* and this mimicked an increase in both plasmatocyte and crystal cell differentiation (Fig. S6A-F) observed upon depletion of Bendless or abrogation of K63 ubiquitination, which also underscores the role of Dishevelled in the LG hematopoietic progenitors. Overall, these results imply that Bendless-mediated K63 ubiquitination is important for the stability and function of Dsh in the LG, failure of which results in inactive Wingless signalling and aberrant blood cell differentiation.

**Figure 4:**
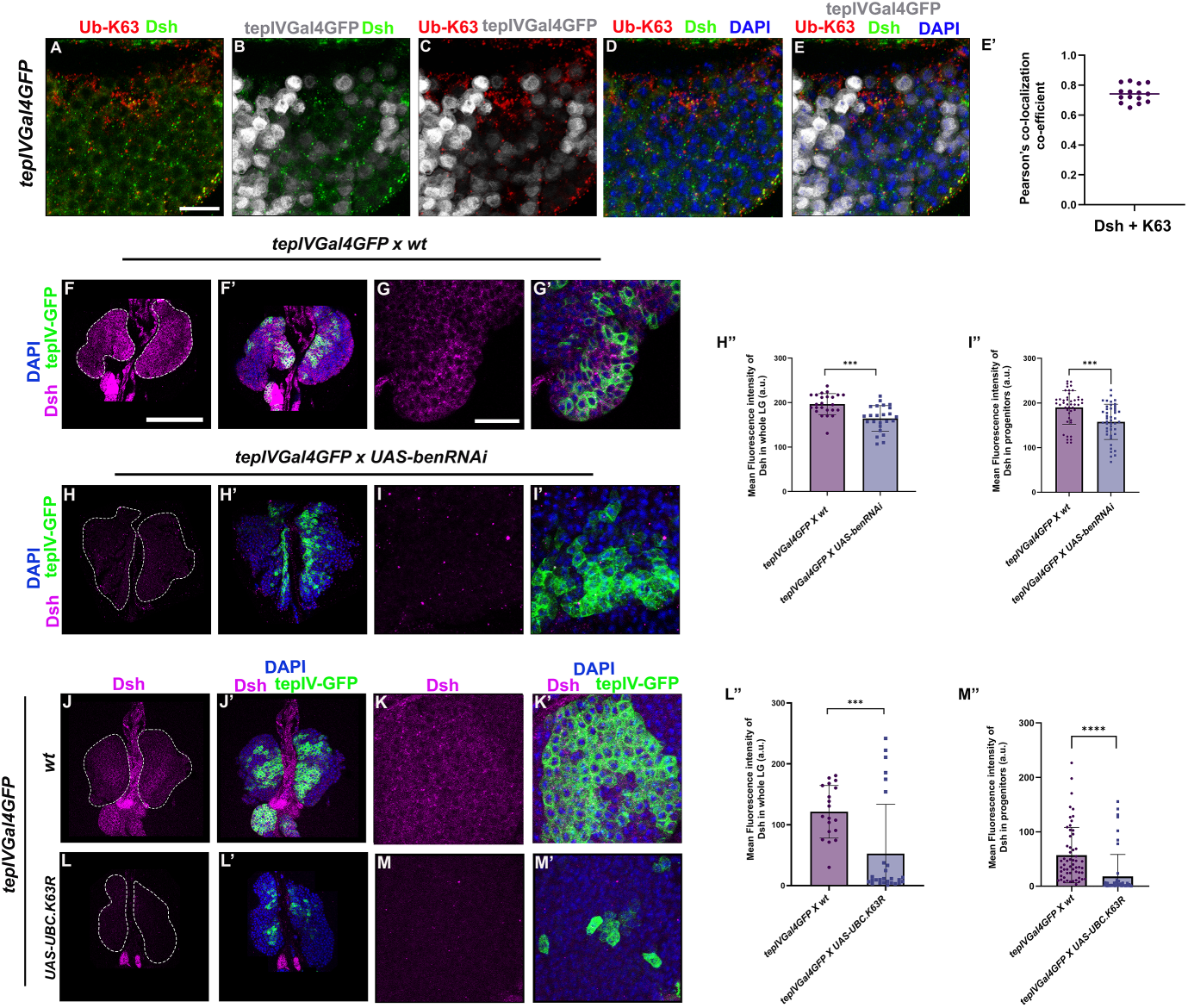
K63-linked ubiquitination regulates Dishevelled in the *Drosophila* LG. Dsh (green) is expressed in the core progenitors marked by tepIVGal4-GFP (grey) and co-localizes with Ub-K63 (red) (A-E). Corresponding Dsh and Ub-K63 co-localization is represented graphically as Pearson’s co-localization coefficient (E’). Dsh expression (magenta) upon *tepIVGal4-*mediated expression of *UAS-benRNAi* (H-I’) or a non-functional ubiquitin mutant, Ubc.K63R (L-M), as compared to their respective wildtype controls (F-G, J-K). High magnification images of Dsh expression in tepIV-GFP positive progenitors upon knockdown of Bendless or upon expression of a non-functional Ubiquitin mutant - Ubc.K63R in tepIV positive progenitors as compared to respective controls (G-I’, K-M’). Corresponding Dsh expression levels are represented as mean fluorescence intensity in the whole primary LG lobe or in progenitors (in arbitrary units; H’’, I’’, L’’, and M’’). Nuclei are stained with DAPI (Blue). GFP (Green) is driven by *tepIV-Gal4*. Scale Bar: 50µm (F-F’, H-H’, J-J’, L-L’), 30µm (A-E, G-G’, I-I’, K-K’, M-M’). Statistical analysis was performed using Student’s t-test with Welch’s correction, and the Pearson correlation coefficient measures the co-localization between Dsh and K63. Each data point represents data from a single LG lobe. P-values denoted are as follows: **** (P<0.0001) and *** (P<0.001).

### Loss of Bendless or abrogation of K63 ubiquitination impairs Wingless signalling output in the hematopoietic progenitors

Our observations related to Dsh in the LG upon Bendless depletion or abrogation of K63 ubiquitination prompted us to examine the status of Wingless signalling in the LG. As shown earlier, we assessed for Wingless ligand and the Frizzled2 receptor as signalling readout for active Wingless signalling (24). This was also shown earlier, where expression of Wg and Fz2, both were shown to be transcriptionally regulated by the Wg pathway itself (57). Notably, both Ben knockdown or expression of UbK63R mutant in tepIV positive progenitors using *tepIV-Gal4* driver, resulted in a significant reduction of both Wg and Fz2 expression levels in the LG, consistent with attenuated Wg signaling activity (Fig. 5A-H’’). Based on these observations, we next asked whether phosphorylated β-catenin (*Drosophila* Armadillo), which is stabilized and translocates to the nucleus when the destruction complex is inhibited, is affected upon Ben depletion or K63 abrogation. We probed this using the pY489-β-cat antibody, which was used in an earlier study where they showed that nuclear pY489-β-cat was observed in the distal region of the medullary zone and in intermediate progenitors (45). Here, we also find that some of the tepIV-GFP cells driven by *tepIV-Gal4* have the presence of nuclear β-cat which decreases dramatically upon tepIV-Gal4-mediated depletion of Bendless or upon abrogation of K63 ubiquitination (Fig. 5I-N’’). These results indicate that Bendless-mediated K63 ubiquitination is important for positively regulating Wingless signalling in the progenitors via Dishevelled.

**Figure 5:**
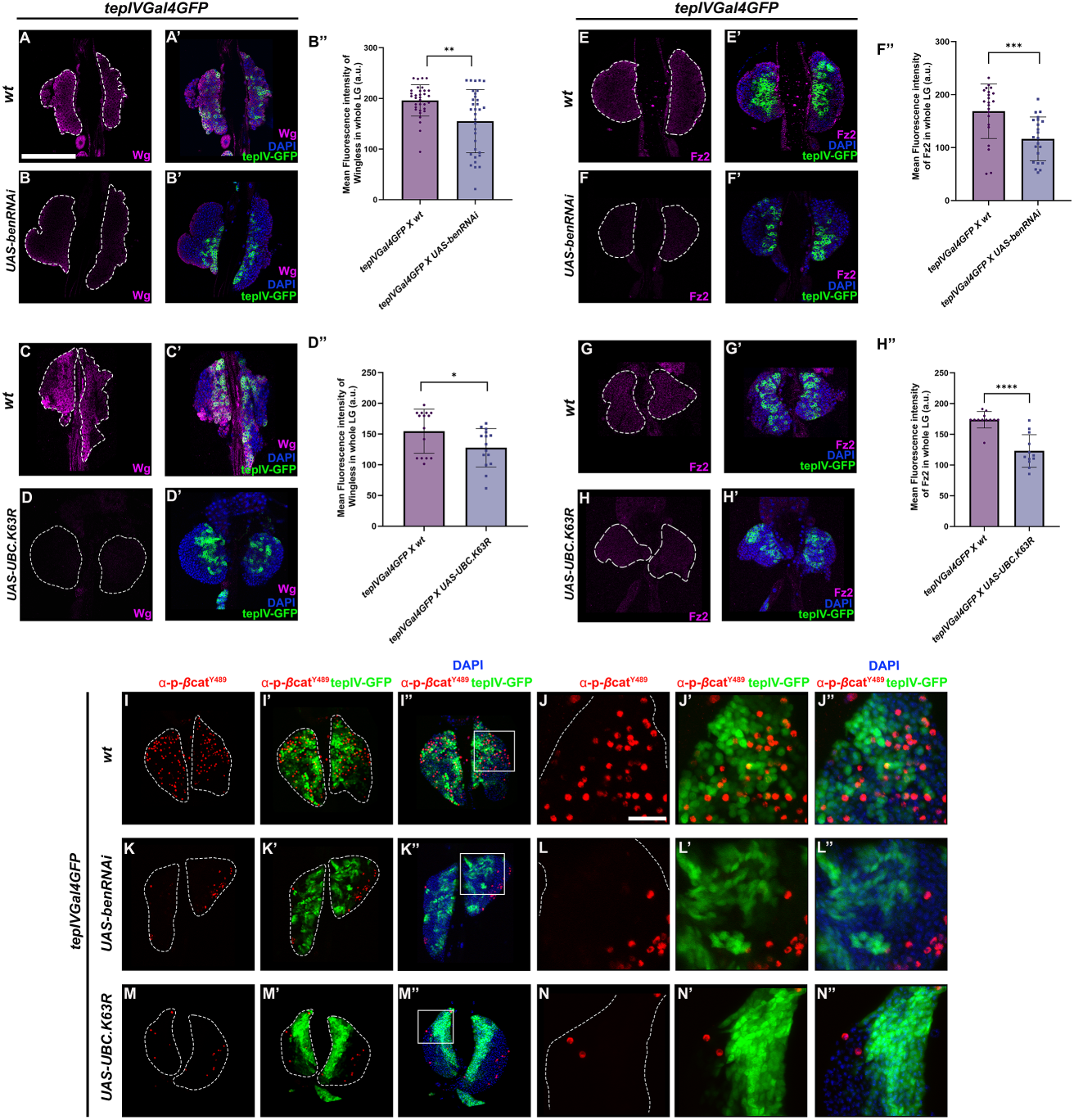
K63 ubiquitination maintains Wg signaling output in LG hematopoietic progenitors. Wg and DFz2 expression (magenta) in LGs bearing *tepIV-Gal4* mediated core progenitor-specific Ben knockdown (B-B’, F-F’) or expression of a non-functional ubiquitin mutant, Ubc.K63R (D-D’, H-H’) as compared to the wild type control (A-A’, E-E’, C-C’, G-G’). Wg and Fz2 levels in the LG lobes are represented as mean fluorescence intensity (in arbitrary units; B’’, D’’, F’’, H’’). Phospho Y489-specific anti-β-catenin antibody (anti-pY489-β-catenin) detecting active nuclear form of β-catenin (red) in LG having *tepIV-Gal4* mediated core progenitor-specific Ben knockdown (K-K’’) or expression of a non functional Ubiquitin mutant - UAS-UBC.K63R (M-M’’) as compared to control (I-I’’). High magnification images (J-N’’) of boxed regions in I’’-M’’. Nuclei are stained with DAPI (Blue). GFP (Green) is driven by *tepIV-Gal4*. Scale Bar: 50µm (A-I’’, K-K’’, M-M’’), 30µm (J-J’’, L-L’’, N-N’’). Statistical analysis was performed using Student’s t-test with Welch’s correction. Each data point represents data from a single LG lobe. P-values denoted are as follows: **** (P<0.0001), *** (P<0.001), ** (P<0.01), and * (P<0.1).

### Excess of Bendless or over-activation of K63 ubiquitination in the hematopoietic progenitors leads to JNK pathway driven lamellocyte production

Previous work reported that Bendless (Ben) acts as a key regulator of the JNK-signalling pathway, acting as an essential upstream component of the Eiger (Egr)-JNK signaling cascade before dTRAF2 to ensure proper JNK activation (7). Additionally, studies have shown that the JNK pathway contributes to lamellocyte differentiation, a specialized hemocyte lineage that is produced upon wasp infestation in *Drosophila* (58–61). From our previous observations, Ben knockdown in progenitors led to an increase in plasmatocyte and crystal cell differentiation. However, Ben over-expression in the tep-IV positive progenitors results in β-integrin (Mys) positive lamellocyte production as compared to the wild type control (Fig. 6A-D, E). Extending these observations, overexpression of the Bendless-associated K63-ubiquitination machinery components—Effete, a highly conserved Class I E2 ubiquitin-conjugating enzyme in *Drosophila*, and Traf6, an E3 ligase—in the tepIV progenitors also results in lamellocyte induction (Fig. 6A-B, F-I, E). We then over-activated K63 ubiquitination by using *UAS-UBC.K63,* where all lysines of ubiquitin except K63 are replaced with arginine. Expression of this construct in the tepIV positive progenitors promotes the formation of K63 linkages (Fig. S5K-O). Expression of *UAS-UBC.K63* in the tepIV region also led to an induction of lamellocytes (Fig. 6J-K, E). Since the JNK pathway is an important determinant of lamellocyte production, we wanted to investigate if Bendless positively regulates the JNK pathway. In order to investigate the activation of the JNK signaling cascade upon Ben overexpression, we examined the expression of several well-characterized JNK pathway target genes by qRT-PCR. Using the pan-hemocyte *Hml-GAL4* driver, we overexpressed Ben in all the entire hemocyte population and probed for the target genes. We detected significant upregulation of *mmp1*, a matrix metalloproteinase and direct transcriptional target of JNK (62), *kayak (kay)*, the *Drosophila* Fos homolog functioning as a downstream effector of JNK signaling (63–64), and *puckered (puc)*, a dual-specificity phosphatase that acts as a negative feedback regulator of the pathway (65) (Fig. 6L–N). In addition to looking at JNK target genes by qRT-PCR, we also checked Mmp1 expression in the LG which gets upregulated upon JNK activation. We found that Mmp1 is turned on with its levels increasing upon over-expression of Ben (Fig. 6P-T, W) or Traf6 (Fig. 6Q-U, W) or upon over-activation of K63 ubiquitination (Fig. 6R-V, W) in the tepIV positive progenitors as compared to the wild type control (Fig. 6O-S, W) again supporting our previous observations that point towards Ben and K63 ubiquitination being positive modulators of JNK pathway for lamellocyte differentiation. The concurrent elevation of these targets further supports the conclusion that Ben overexpression activates the JNK signaling axis in the LG. These findings provide strong molecular and genetic evidence that Ben promotes lamellocyte formation via K63-linked polyubiquitination-dependent activation of the JNK pathway, and highlight the role of this specific ubiquitin linkage type in modulating stress-responsive signaling during hematopoiesis.

**Figure 6:**
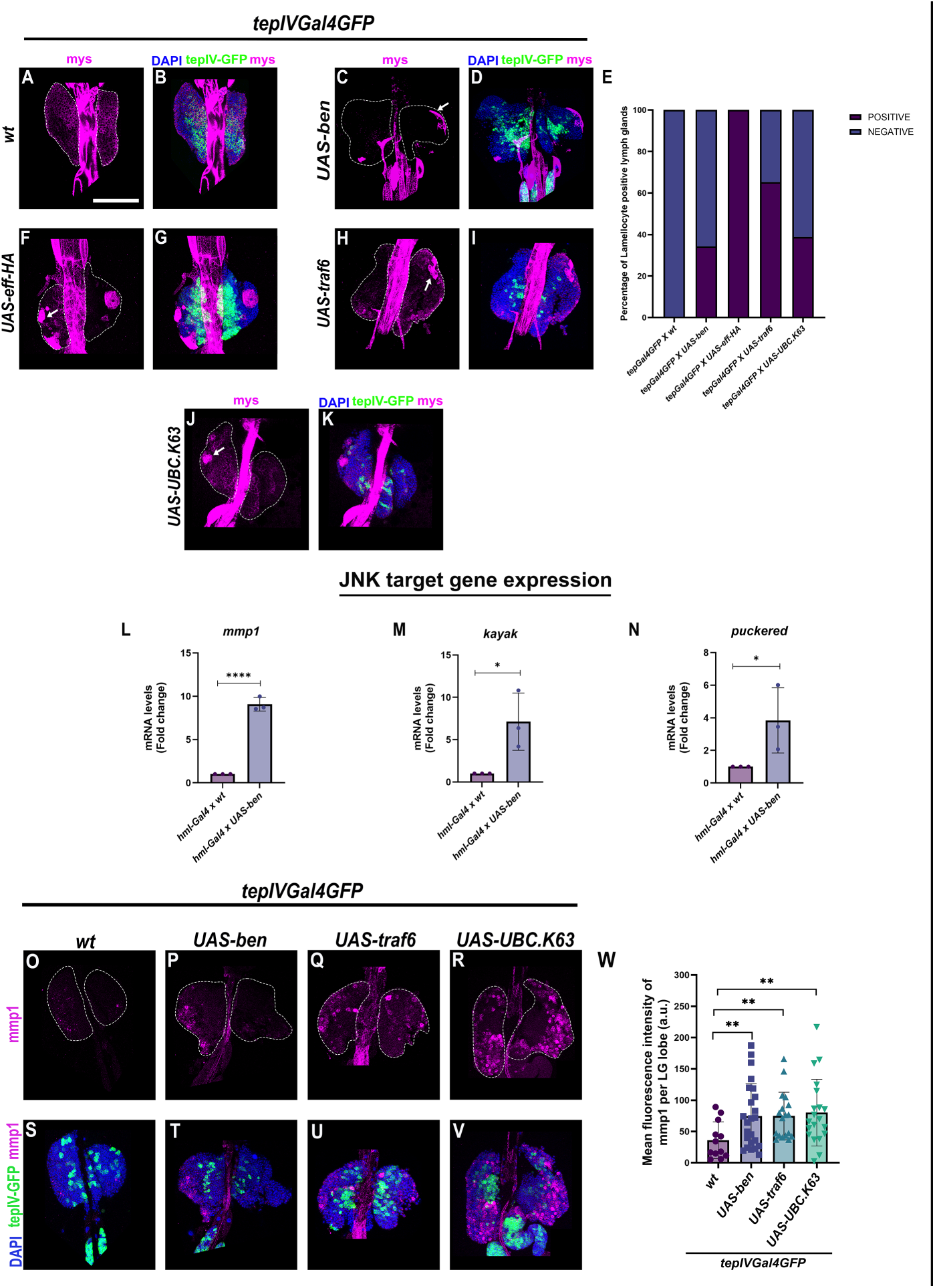
Over-activation of K63 ubiquitination or Bendless over-expression in hematopoietic progenitors leads to lamellocyte production via the JNK pathway. Lamellocyte differentiation marked by Myospheroid (Mys) (magenta) upon *tepIVGal4* mediated expression of *UAS-ben* (C-D) or *UAS-eff-HA* (F-G) or *UAS-traf6* (H-I) or *UAS-UBC.K63* (J-K) as compared to *wildtype* control (A-B). Percentage of lamellocyte-positive lymph glands represented graphically (E). Relative *mmp1, kayak,* or *puckered* mRNA transcript levels determined by qRT-PCR upon pan-hemocyte-specific (using *hmlGal4*) expression of *UAS-ben* as compared to *wildtype* control (L-N). Mmp1 expression (magenta) in the LG upon *tepIVGal4* mediated expression of *UAS-ben* (P, T) or *UAS-traf6* (Q, U) or *UAS-UBC.K63* (R, V) as compared to wildtype control (O, S). Corresponding mean fluorescence intensity of mmp1 levels in the LG lobe upon *tepIVGal4* mediated expression of *UAS-ben* or *UAS-traf6* or *UAS-UBC.K63* as compared to wildtype control (W). Statistical analysis for qRT-PCR was performed using One-way ANOVA (Dunnett test) for comparison of all test genotypes with the wild-type control. GFP expression (green) is driven by *tepIVGal4*. Nuclei are stained with DAPI (Blue). Scale Bar: 50µm (A-D, F-K, O-V). Statistical analysis was performed using Student’s t-test with Welch’s correction. Each data point represents data from a single LG lobe. P-values denoted are as follows: **** (P<0.0001), ** (P<0.01), and * (P<0.1).

### Genetic epistasis analysis substantiates that Bendless mediated K63 ubiquitination regulates progenitor maintenance and lamellocyte differentiation via Wingless and JNK pathway respectively

Since Bendless depletion or abrogation of K63 ubiquitination in tepIV progenitors affected Dishevelled in the LG thereby impacting Wingless signalling activity as evidenced by Wg, Fz2 expression and nuclear β-catenin localization we wanted to test if downstream activation of Wingless pathway is capable of genetically rescuing the aberrant plasmatocyte and crystal cell differentiation phenotypes in the LG. We utilized the use of knockdown of Shaggy (sgg), the *Drosophila* homolog of GSK3β, a key negative regulator within the β-catenin destruction complex. GSK3β phosphorylates Armadillo (*Drosophila* β-catenin), targeting it for degradation, and its loss results in stabilization and nuclear translocation of Armadillo, thus enhancing Wnt signaling (66–69). We reasoned if the Ben depletion induced phenotype is mediated via Wingless signaling, then co-depletion of Sgg in this genetic background should rescue the excess differentiation phenotypes. Our results show that depletion of Sgg in the genetic background of Ben knockdown in tepIV progenitors resulted in a signficant genetic rescue of both P1 positive plasmatocyte and Hnt positive crystal cell differentiation (Fig. 7C-F, G-H) as compared to the Ben depletion control (Fig. 7B-E, G-H) whereas depletion of Sgg alone led to hemocyte differentiation very similar to a wild type (Fig. 7A-D, G-H). These results provide genetic evidence for Bendless mediated K63 ubiquitination being upstream regulators of Wingless signalling in the LG.

**Figure 7:**
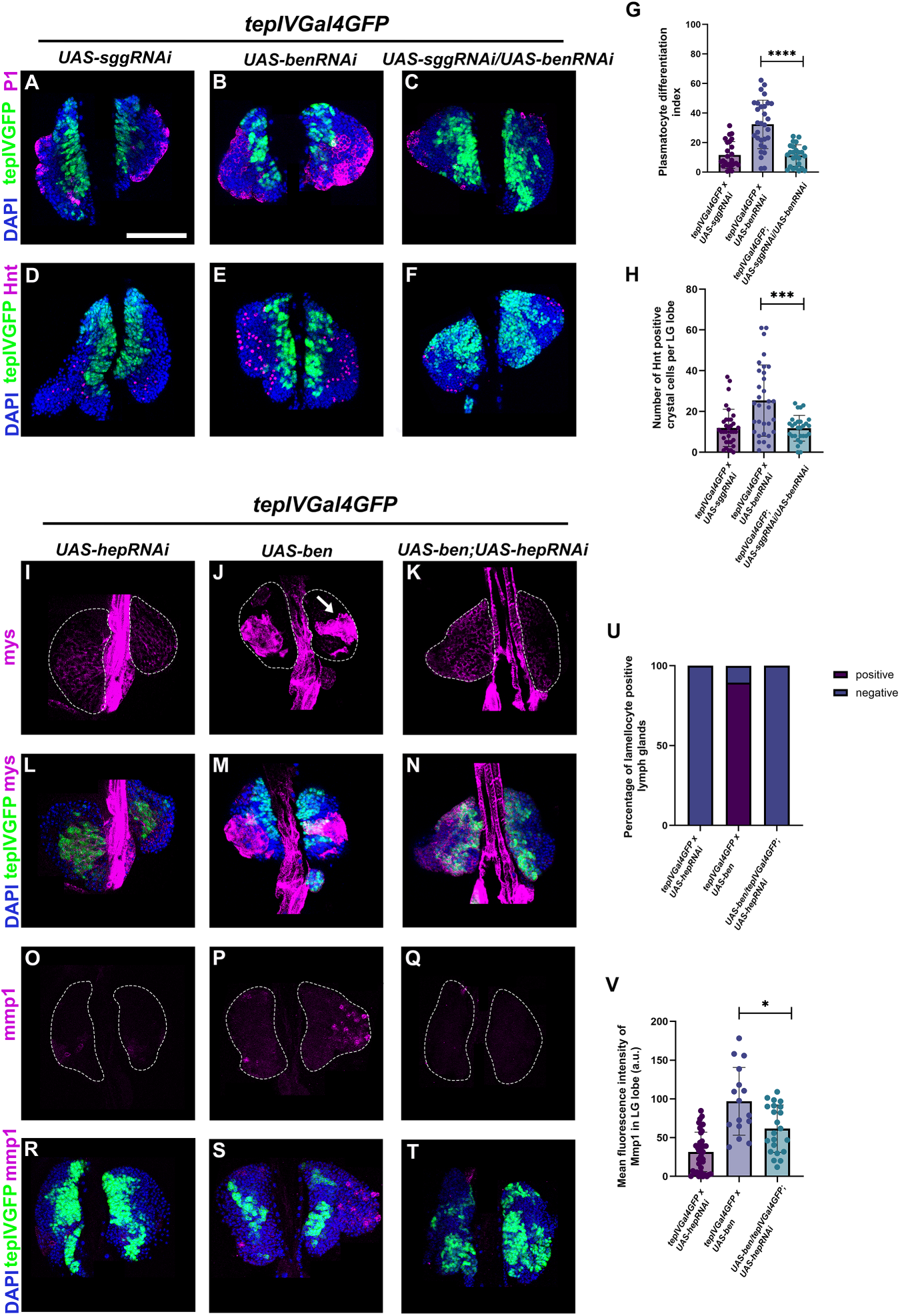
Genetic epistasis analysis shows that Wingless pathway activation or JNK pathway inactivation can rescue Bendless depletion and over-expression LG phenotypes, respectively. Plasmatocyte differentiation marked by P1 (magenta) or crystal cell differentiation marked by Hindsight (Hnt, magenta) upon *tepIVGal4* mediated depletion of Shaggy (Sgg) viz. *UAS-sggRNAi* (C, F) in the *bendless* knockdown genetic background (via *UAS-benRNAi*) as compared to the Ben or Sgg alone knockdown controls (A-B, D-E). Graphical representation of plasmatocyte differentiation index or total number of crystal cells upon *tepIVGal4-*mediated depletion of Sgg in a Ben knockdown genetic background as compared to wildtype control (G-H). Lamellocyte differentiation marked by Myospheroid (Mys, magenta) or JNK target Mmp1 expression (magenta) upon *tepIVGal4* mediated depletion of hemipterous (Hep) using *UAS-hepRNAi* in the Bendless over-expression genetic background (via *UAS-ben*) (K, N, Q, T) as compared to Hep knockdown or Ben over-expression control (I-J, L-M, O-P, R-S). Graphical representation of percentage of lamellocyte positive lymph glands (U) or mean fluorescence intensity of Mmp1 upon *tepIVGal4* mediated expression of *UAS-hepRNAi* in a Ben overexpression background as compared to Hep knockdown or Ben overexpression control (U-V). GFP expression (green) is driven by *tepIVGal4*. Nuclei are stained with DAPI (Blue). Scale Bar: 50µm (A-F, I-T). Statistical analysis was performed using Student’s t-test with Welch’s correction. Each data point represents data from a single LG lobe. P-values denoted are as follows: **** (P<0.0001), *** (P<0.001), and * (P<0.1).

Now, in addition to the role of Ben in regulating Wingless signalling we wanted to delineate its role in the JNK pathway activation. Our findings showed that an excess of Ben or over-activation of K63 ubiquitination in the hematopoietic progenitors resulted in lamellocyte induction and we found that key targets of the JNK pathway are turned on upon Ben over-expression. In order to genetically delineate that Ben indeed acts upstream in the pathway to control lamellocyte differentiation, we decided to deplete a downstream component of the JNK pathway named Hemipterous (Hep kinase) in the genetic background of Ben over-expression in the tepIV positive hematopoietic progenitors. Depletion of Hep in Ben over-expression genetic background in hematopoietic progenitors rescued the lamellocyte differentiation phenotype where none of the LGs were lamellocyte positive (Fig. 7K-N, U) as compared to the Ben over-expression genotype where lamellocytes were present (Fig. 7J-M, U) and Hep alone depletion control where no lamellocytes were observed (Fig. 7I-L, U). We also examined the JNK target, Mmp1 in the LG upon downstream inactivation of JNK in Ben over-expression background in the blood progenitors. We observe that Hep depletion in Ben over-expression genetic background led to a decrease in Mmp1 levels (Fig. 7Q-T, V) as compared to the Ben over-expression genetic background (Fig. 7P-S, V) and Hep depletion alone control where Mmp1 levels are low (Fig. 7O-R, V). The genetic epistasis data shows that Bendless is an upstream regulator of JNK pathway and is a positive regulator as excess Bendless leads to an increase in JNK pathway activation.

Taken together, our data suggests that optimal levels of Bendless mediated K63 ubiquitination are important for regulating blood cell homeostasis in the *Drosophila* LG where both loss or excess of Bendless can lead to a disruption of blood cell homeostasis (Fig. 8). We propose that a molecule like Bendless that controls K63 ubiquitination can potentially act as an internode connecting multiple signalling pathways and enabling cross-talk and facilitating a calibrated response during developmental or disease conditions. Our findings underscore a novel link between ubiquitin-mediated signaling regulation and blood cell differentiation, expanding the understanding of how non-degradative ubiquitination by Bendless influences *Drosophila* hematopoiesis.

**Figure 8:**
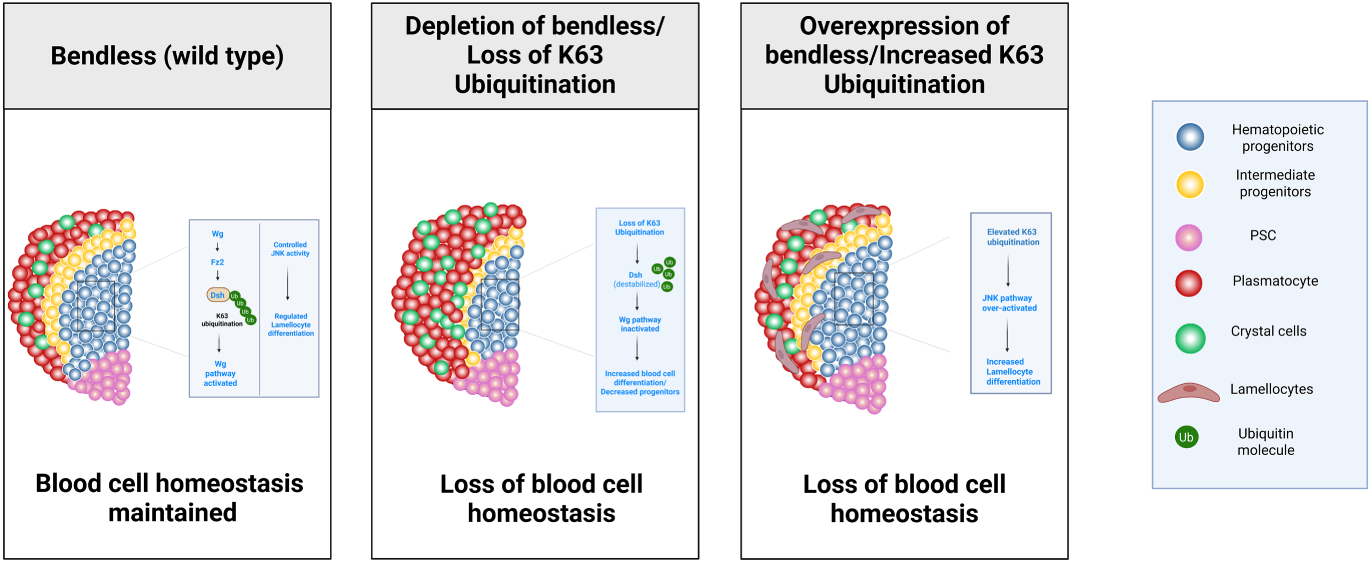
Bendless-Dependent K63 Ubiquitination maintains Blood Cell Homeostasis in *Drosophila.* A mechanistic model depicting the role of Bendless-mediated K63 ubiquitination in regulating hematopoietic progenitor maintenance and blood cell differentiation. Optimal levels of Bendless and K63 ubiquitination are critical for maintaining blood cell homeostasis. Abrogation of K63 ubiquitination or Bendless depletion in hematopoietic progenitors leads to inactive Wingless signalling, resulting in loss of progenitor maintenance and increased blood cell differentiation. Over-activation of K63 ubiquitination or excess Bendless in hematopoietic progenitors results in activation of the JNK pathway mediated lamellocyte differentiation. This dual regulation maintains progenitor identity while controlling immune-driven hematopoiesis, offering mechanistic insight into how ubiquitin-mediated signaling shapes the hematopoietic microenvironment. Created in BioRender. Khadilkar, R. (2025) https://BioRender.com/f81et3q

## Discussion

Polyubiquitylation of proteins through distinct lysine residues on ubiquitin generates a versatile code that regulates a wide range of cellular functions. Among these, K63-linked polyubiquitination has emerged as a key non-proteolytic modification involved in regulating signaling cascades, DNA repair, vesicular trafficking, and immune responses (3–4). Although its role has been extensively studied in immune receptor signaling (3,72), the *in-vivo* role of K63 ubiquitination in regulating developmental hematopoiesis remains poorly understood. In mammals, conditional Ubc13 knockout models display profound hematopoietic abnormalities, including depletion of progenitor pool and failure of multilineage differentiation (13), indicating a cell-intrinsic role for K63-linked ubiquitin signaling in stem and progenitor maintenance.

In *Drosophila*, larval LG hematopoiesis is governed by a network of evolutionarily conserved developmental pathways, including Wingless (Wg/Wnt), Hedgehog (Hh), Dpp/BMP, Notch, JAK/STAT, and Hippo, which coordinate progenitor maintenance, proliferation, and differentiation (16, 18, 20, 24, 31, 34, 70, 71). These pathways function in a spatiotemporally regulated manner within lymph gland compartments such as the Posterior Signaling Center (PSC) and Medullary Zone (MZ) to balance progenitor maintenance and blood cell production for immune readiness or stress-induced hematopoiesis. Since there are multiple signalling pathways that function in tandem and consist of components having a cross-talk with each other, understanding their fine modulation becomes important. Here, we delineate the role of K63 polyubiquitination in controlling signalling pathways that regulate LG developmental hematopoiesis.

Our study focuses on understanding the role of an E2 conjugating enzyme involved in K63 ubiquitination named Bendless in LG hematopoiesis. Bendless is expressed in the LG hematopoietic progenitors that show active K63-linked polyubiquitin chains indicating an important role for K63 ubiquitination in this developmental zone. To assess the physiological relevance of Ben, we analyzed whole-animal mutants for *ben* and Traf6, an E3 ligase that functions with Ben (73–75). Both *ben* or *traf6* mutant alleles exhibited a significant reduction in Antennapedia (Antp) positive PSC niche cells, a phenotype reminiscent of defects in insulin/TOR signaling, which has been shown to regulate PSC cell number and niche maintenance (26). This suggests that K63 ubiquitination could be modulating signals that regulate hematopoietic niche size which needs further exploration. In addition to the niche phenotype, these mutants displayed increased plasmatocyte or crystal cell differentiation indicating an important role of these molecules in controlling blood cell differentiation. We also found lamellocytes in these mutants which suggests that signalling pathways like Toll, Imd or JAK/STAT that have been previously linked to lamellocyte production (58, 76, 77–84) could be potentially deregulated in these mutants giving rise to lamellocytes. Since the mutant alleles are whole animal mutants of Bendless, it is quite likely that multiple signalling pathways might be affected at a systemic level which could impact the hematopoietic organ leading to increased blood cell differentiation given that the organ is very sensitive to systemic cues (26).

Since Bendless is expressed in the LG hematopoietic progenitors that display active K63 ubiquitination and the mutant alleles showed an abrogated blood cell homeostasis phenotype, we perturbed Bendless expression or K63 ubiquitination in the progenitors. Localized depletion of Bendless or abrogation of K63 ubiquitination in the progenitors led to a significant decrease in tepIV positive core progenitors with a concomitant increase in plasmatocyte and crystal cell differentiation. Interestingly, depletion of associated molecules that work with Bendless like Uev1a, Effete or Traf6 also led to a similar increased differentiation phenotype indicating an important role for Bendless mediated K63 ubiquitination in regulating the balance between tepIV positive core progenitors and differentiated cells. Multiple developmental signalling pathways control this fine balance however we hypothesized that Wnt (Wingless) signalling might be affected due to abrogation of K63 ubiquitination. Now, there have been conflicting reports in the past on how K63 ubiquitination could be potentially regulating Wnt. In the context of human cylindromas, loss of CYLD - a deubiquitinase results in enhanced K63 ubiquitination of Dishevelled (Dvl) an upstream component of Wnt signalling pathway that gets stabilized leading to activated Wnt signalling (53). On the other hand, a conflicting report shows that loss of Ubc13 results in accumulation of β-catenin and increased expression of Wnt target genes resulting in impaired hematopoiesis (13). We examined the possibility of Dsh being K63 ubiquitinated in the LG. We studied the subcellular localization of Dsh and K63-linked ubiquitin chains in lymph gland progenitors and observed a strong co-localization, indicating a potential functional interaction in the progenitors. Loss of Bendless or abrogation of K63 ubiquitination activity affected the stability and localization of Dishevelled in the LG which in turn impacted Wingless signalling activity in the progenitors resulting in loss of progenitor maintenance and increased differentiation. There is increased blood cell differentiation upon Dishevelled knockdown in hematopoietic progenitors corroborating our findings on the importance of Dishevelled in regulating Wingless pathway in LG blood cell progenitors. Canonical activation of Wingless pathway by depletion of Shaggy (Sgg) from hematopoietic progenitors in Bendless knockdown genetic background restored the blood cell differentiation phenotype. Now, it remains to be tested if depletion of Bendless or abrogation of K63 ubiquitination has a similar effect on other progenitor populations in the LG namely the distal and intermediate progenitors. There is a possibility that Bendless mediated K63 ubiquitination could be regulating signalling components from other key pathways like JAK/STAT (31, 85), Hedgehog (18, 20), Dpp signalling (20, 86) that are active in the progenitors and would need further investigation.

While the depletion of Bendless or abrogation of K63 ubiquitination leads to progenitor exhaustion owing to inactive Wingless signalling, our observations indicate that excess of Bendless or over-activation of K63 ubiquitination leads to increased lamellocyte differentiation. Now there are multiple signalling pathways like JNK, EGFR, JAK/STAT, Toll pathways that are linked to have a role in lamellocyte differentiation upon wasp infestation (16, 87–89). We specifically examined if Bendless mediated K63 ubiquitination modulates JNK pathway in the context of lamellocyte differentiation as K63 ubiquitination has been reported to modulate upstream components of the JNK pathway wherein K63 poly-ubiquitin chains function as scaffolds facilitating protein-protein interactions enabling the activation of kinases in the JNK cascade (90–92). It was also reported that an upstream adaptor of the TNF receptor pathway, TRAF2 is K63 ubiquitinated leading to activation of downstream kinases. This modification is largely catalyzed by Ubc13, the mammalian counterpart of Bendless (90). Bendless has been shown to be associated with the JNK signaling pathway through its role as an E2 ubiquitin-conjugating enzyme that mediates K63-linked polyubiquitination of dTRAF2, a critical adaptor in the TNF/Eiger signaling cascade (7). This post-translational modification facilitates the activation of downstream kinases such as TAK1 and Hemipterous (Hep), ultimately leading to the activation of JNK (Basket) and AP-1-dependent transcriptional responses involved in processes such as cell death, migration, and development (93). This prompted us to look into the K63 ubiquitination connection with lamellocyte differentiation via JNK. Our results show that JNK downstream targets are turned on when Bendless is over-expressed or K63 ubiquitination is over-activated and the lamellocyte differentiation phenotype is genetically rescued upon depletion of a downstream kinase in JNK pathway named Hemipterous (Hep) in the genetic background of Bendless over-expression. While we have investigated the connection of JNK in this context, there is a strong possibility that NF-KB signalling pathways that play an important role in lamellocyte production could be modulated by Bendless as well which warrants further investigation. It is also notable that K63-linked ubiquitination is a well-known modulator of NF-κB and JNK pathway components in mammalian immune signaling, especially through TRAF6, TAK1, and TAB2/3 complexes (3, 94), further strengthening the conservation of this regulatory axis. We speculate that wasp infestation could be a trigger for formation and linkage of K63 poly-ubiquitin chains on target molecules that are then capable of sensing wasp induced stress to switch over to an emergency hematopoiesis mode for lamellocyte differentiation to counter the stress.

Given that Wg signaling maintains blood progenitors and prevents their aberrant differentiation (24), and JNK signaling has dual roles in promoting immune activation and modulating apoptosis and differentiation under stress (95–97), our results suggest that Bendless may integrate these pathways to preserve lymph gland homeostasis. In addition to Wg and JNK, it remains to be explored whether Notch, which drives crystal cell fate (98), or Hippo signaling, implicated in progenitor proliferation and immune activation (71, 99), are influenced by ubiquitin-mediated control. The possible regulation of these pathways by Ben or associated K63-linked machinery could provide deeper insights into the post-translational crosstalk underlying hematopoietic balance.

We speculate that Bendless mediated K63 ubiquitination could be acting as an inter-node that can potentially help in switching over cellular states based on either local or systemic cues facilitating reinforcement of not only developmental homeostasis but also acting as a turn-on switch for stress or emergency induced changes in homeostasis. Taken together, our findings reveal a dual role for Bendless in sustaining progenitor identity and regulating immune-activated hematopoiesis, providing pertinent mechanistic insights into how ubiquitin-mediated signaling shapes the hematopoietic microenvironment.

## Materials & Methods

### *Drosophila* Genetics and husbandry

All the *Drosophila* stocks and crosses were maintained at 25°C, in a standard cornmeal diet containing corn starch & sugar as carbon source, malt extract containing trace number of vitamins and minerals, yeast extract as protein source and agar as solidifying agent. *Canton-S* was used as *wildtype* control. Tissue specific *Gal4* promoter line was used to drive the expression of *UAS* responder genes. Respective *Gal4* parent stocks crossed to wild type strain as *Canton-S* were used as controls wherever appropriate. *Gal4* driver lines used were *tepIVGal4GFP* on Chr. II*, domeGal4GFP* on X*, e33cGal4* on Chr. III (gifted by Dr. Maneesha Inamdar, JNCASR & DBT-InStem). The *UAS*-transgene lines used were *UAS-benRNAi* (Chr III, RRID:BDSC_28721), *UAS-ben* (Chr. II, Gift from Dr. Manish Jaiswal, TIFR Hyderabad), *UAS-effeteRNAi* (Chr. III, Gift from Dr. Sonal Nagarkar Jaiswal, CSIR-CCMB), *UAS-traf6RNAi* (Chr. II, RRID:VDRC_110266), *traf6^ex1^* (Chr. II, RRID:BDSC_82149), *UAS-traf6* (Chr. II, RRID:BDSC_82150), *UAS-sggRNAi* (Chr. III, RRID:BDSC_31308), *UAS uev1aRNAi* (Chr. II, RRID:BDSC_66947) *UAS-dshRNAi* (Chr. III, RRID:BDSC_31307) *UAS-hepRNAi* (Chr. III, RRID:BDSC_28710), *UAS-eff.HA* (Chr. II, RRID:BDSC_26691), w*; *P{UAS-UBC.K63}4/CyO* (Chr. II, RRID:BDSC_32054), w*; *P{UAS-UBC.K63R}5/CyO* (Chr. II, RRID:BDSC_32053), *y^1 w*^ ben^B P^{neoFRT}19A/FM7c, P{GAL4^-^ Kr.C}DC1, P{UAS-GFP.S65T}DC5, sn^+^* (Chr. X, RRID:BDSC_57058), *y^1 w*^ ben^A P^{neoFRT}19A/FM7c, P{GAL4-Kr.C}DC1, P{UAS-GFP.S65T}DC5, s^n+^* (Chr. X, RRID:BDSC_57057), *w^1118 b^e^n1/^C*(*1*)*A, ^y1^* (Chr. X, RRID:BDSC_9565).

### *Drosophila* genetic crosses

#### Multi-generation/epistasis experiments

To determine if activation of Wingless signalling can genetically rescue the increased hemocyte differentiation phenotype upon Bendless depletion in tepIV positive progenitors, a knockdown line of Shaggy (Sgg) was placed in the genetic background of Bendless depletion in tepIV positive progenitors (*tepIV-Gal4*) Resultant genotype: *tepIVGAL4GFP/tepIVGAL4GFP, UAS-sggRNAi/UAS-benRNAi*

To determine if JNK pathway in-activation can genetically rescue the lamellocyte differentiation phenotype upon Bendless over-expression using *tepIV-Gal4*, hemipterous (hep) was depleted in Bendless over-expression genetic background using *tepIV-Gal4*: Resultant genotype: *tepIVGAL4GFP/UAS-ben; UAS-hepRNAi*/*UAS-hepRNAi*

#### Antibodies

Antibodies used were mouse-raised anti-P1 (1:100, kind gift from Dr. Istvan Ando), mouse-raised anti-Hindsight (1:25, 1G9 – DSHB; RRID:Ab_528278), mouse-raised anti-Antp (1:25, 8C11-DSHB; RRID:Ab_528083), mouse-raised anti-γ2AX (1:200, UNC93-5.2.1 - DSHB; RRID:Ab_2618077), mouse-raised anti-Myospheroid (1:25, 6G11 - DSHB; RRID: Ab_528310), mouse-raised anti-Mmp1 (1:25, 5H7B11-s-DSHB; RRID: AB_579779), mouse-raised anti-Mmp1 (1:25, 3B8D12-s-DSHB; RRID: AB_579781), mouse-raised anti-Mmp1 (1:25, 3A6B4-s-DSHB; RRID: AB_579780), mouse-raised anti-Wg (1:25, 4D4-s-DSHB; RRID: AB_528512), mouse-raised anti-Fz2 (1:25, 12A7-s-DSHB; RRID: AB_528257), rabbit-raised anti Ub-K63 (1:25, JM09-67-Invitrogen; RRID: AB_2809850), rabbit-raised anti-Bendless (1:50 Gift from Dr. Manish Jaiswal, TIFR Hyderabad) for immunofluorescence based experiments. Normal Goat Serum (HIMEDIA, RM10701) was used as the blocking agent. Alexa-Fluor 568 conjugated secondary antibodies – Goat-raised anti-Mouse 568 (1:400, Invitrogen, RRID: Ab_144696), Goat-raised anti-Mouse 633 (1:400, Invitrogen, RRID: AB_2535719) and Goat-raised anti-rabbit 568 (1:400, Invitrogen, RRID: AB_10563566) were used for immunofluorescence-based experiments.

#### Lymph Gland Dissection, Immunohistochemistry & Mounting

For the dissection of lymph glands, wandering larvae in their late third instar were used. Dissections were carried out in phosphate buffer saline (PBS), fixed for 20 minutes in 4% paraformaldehyde, and then washed three times for five minutes each in PBS containing 0.3% Triton-X (PBST). After blocking the samples with 20% normal goat serum for 20 minutes at room temperature, the samples were incubated with primary antibodies for the entire night at 4°C. PBST washes, blocking, and treatment with the proper Alexa-Fluor conjugated secondary antibody incubation for two hours at room temperature followed. Three 5-minute PBST washes were performed after this. After that, the LGs were put in Vectashield mounting medium with DAPI (Vector Laboratories, RRID: AB_2336790).

#### Image acquisition and analysis of various Lymph Gland parameters

Confocal images were captured using either a Zeiss LSM 780, Leica SP8 or Nikon Ax confocal microscope. Z projection of the confocal images were used for estimating various lymph gland parameters using ImageJ/Fiji software. Plasmatocyte Differentiation Index was estimated by measuring the percentile of P1 positive area divided by the total area of the primary lobe. The Prohemocyte Index was estimated by measuring the percentile of TepIV-GFP positive area divided by the total area of primary lobe. Freehand selection tool was used for measuring the area of the plasmatocytes or the prohemocytes. For the quantification of Antp and Hnt, the positive signals for respective markers were manually counted using the multi-point tool. The lymph gland quantifications were done for individual primary lymph gland lobes. For estimation of Wg, Fz2, Dsh, Mmp1 levels in the lymph gland using ImageJ/Fiji software (RRID:SCR_003070), the images were acquired at the same intensity settings/parameters and mean fluorescent intensity (represented as arbitrary units) was calculated keeping the threshold value of fluorescent intensity uniform as that of control for all the crosses. Similar protocol was followed for estimating the mean fluorescence intensity of Wg, Fz2, Mmp1 and Dsh in all the crosses, which was calculated keeping the threshold value of fluorescent intensity uniform as that of control for all the crosses. The raw images were processed into .tiff file format (in RGB mode) into single or merge channels as per requirement using ImageJ. These .tiff files were then opened using Adobe Photoshop (Creative Cloud version) and were assembled and stitched as per requirement of the figure panel using a black background on a Photoshop canvas (RGB mode). Final assembled figure panels were saved in TIFF format using LZW compression with a resolution of 600 dpi or more.

#### Pearson’s co-localization co-efficient

Pearson’s co-localization co-efficient was obtained using Co-localization Finder (ImageJ plugin, NIH) by selectively placing the ROI (Region of Interest) tool on the cells in the tepIV-Gal4GFP positive region of the lymph gland.

#### Quantitative Real Time PCR

Third instar larvae (a minimum of 10 larvae) were homogenized and lysed in TRIzol (Ambion – life technologies, Catalogue No.: 11596018). RNA was isolated by pooling all the aqueous layers post-chloroform treatment, followed by RNA isolation according to the manufacturer’s protocol. RNA yield was quantified using Nanodrop. 1μg of mRNA was reverse transcribed using oligo-dT primers (Promega, C110A) and ImProm-II (Promega, A3800). Quantitation of the mRNA transcripts was done using SYBR green chemistry (Thermo Fisher Scientific, Catalogue No.: 4367659) in the Quantstudio 5 RT PCR system (Thermo Fisher Scientific) in quadruplets of 10μl reaction. The data was analyzed using the ΔΔCt method and relative mRNA expression was normalized to *rp49*. Fold change calculations were done in comparison to *wildtype* control. The experiment was done in biological triplicates & statistical analysis was performed using One-way ANOVA (Dunnett) for comparison of all test genotypes with *wildtype* control genotype.

### Statistical analysis

Immunofluorescence based experiments and their analysis were performed on at least 10-15 lymph glands dissected from wandering late third-instar larvae. Statistical analysis was performed using the GraphPad Prism Version 10 software (RRID:SCR_002798). For analysis of statistical significance, each experimental sample was tested with its respective control in a given experimental setup for all the data in each of the figures in order to estimate the P value. P values were determined by using a two-tailed unpaired Student’s t-test with Welch’s correction. Values are mean ± SD, and asterisks denote statistically significant differences (ns denotes not significant, * p < 0.05, ** p < 0.01, *** p < 0.001, **** p < 0.0001). Mutant genotypes were compared to the wildtype controls and the knockdown or over-expression genotypes were compared to their respective parental Gal4 controls that were crossed to wild-type for all the statistical analysis performed. No statistical method was used to pre-determine the sample size and the experiments were not randomized.

### List of Primers

qRT primers are in 5’ to 3’ direction: kayak FP TCCTGGGCATGGGTATTCC kayak RP TGCGTATCGTTGTACGAAGCG mmp1 FP CTTCGCGGGACTGAACATCA mmp1 RP TAGGTGAGGTTCTTCACACGC puckered FP TCCCAATGAGAGCCATCTGC puckered RP GAACCCGTTTTCCGTGCATC

## Acknowledgements

We would like to thank the Digital Imaging Facility (DIF) and the Common Instrumentation Facilities (CIF) at ACTREC for all the support. We thank the Bloomington Drosophila Stock Center, Developmental Studies Hybridoma Bank and the fly community for fly stocks and antibodies. We would like to particularly thank Manish Jaiswal, David Strutt for various fly lines and reagents. We are thankful to the Stem Cell and Tissue Homeostasis lab for useful input and discussions.

## Author contributions

Conceptualization: R.J.K.; Data curation: R.J.K., A.K., Y.S.; Formal analysis: A.K., Y.S., C.P., M.M., R.D., A.S.; Funding acquisition: R.J.K.; Investigation: A.K., Y.S., C.P., M.M., R.D., N.S., A.S.; Methodology: A.K., Y.S., C.P., M.M., R.D.; Project administration: R.J.K.; Resources: R.J.K., M.J.; Supervision: R.J.K.; Validation: A.K., Y.S.; Visualization: R.J.K.; Writing – original draft: A.K., R.J.K.; Writing – review & editing: A.K., R.J.K.,Y.S.

## Funding

This study was funded by Department of Biotechnology, Ministry of Science and Technology, India for the Har Gobind Khorana – Innovative Young Biotechnologist Award (no. BT/13/IYBA/2020/14) to R.J.K., Ramalingaswami Re-entry Fellowship from the Department of Biotechnology, Ministry of Science and Technology, India (BT/RLF/Re-entry/19/2020) to R.J.K., Council of Scientific and Industrial Research (CSIR)-JRF to A.K. This work was also funded by a Basic and Translational Research in Cancer grant (no.1/3(7)/2020/TMC/R&D-II/8823 Dt.30.07.2021), Capacity Building and Development of Novel and Cutting-edge Research Activities (no.1/3(4)/2021/TMC/R&D-II/15063 Dt.15.12.2021) from the Department of Atomic Energy, Government of India.

## Data availability

All relevant data can be found within the article and its supplementary information

## Competing interests

The authors declare no competing or financial interests.

## Notes

### Competing Interest Statement

The authors have declared no competing interest.

