## Supplementary Information for "Bendless-mediated K63 ubiquitination modulates cellular signalling to regulate *Drosophila* hematopoiesis"

**Supplementary Figure legends:**

**Figure S1: *bendless* mutant alleles display lamellocyte-positive lymph glands**

Lamellocyte differentiation marked by Myospheroid (Mys) (magenta) in *ben* mutant alleles ($ben^{1}$, $ben^{A}$, $ben^{B}$) (B-D, F-H) as compared to the wildtype control (A, E). Graphical representation of the percentage of lamellocyte-positive lymph glands in *ben* mutant alleles ($ben^{1}$, $ben^{A}$, $ben^{B}$) as compared to wildtype control (I). Nuclei are stained with DAPI (Blue). Scale Bar: 50µm (A-H). A minimum of N=15 larvae were analyzed for the presence or absence of lamellocytes

**Figure S2: Validation of Bendless knockdown and over-expression lines using Bendless-specific antibody**

Bendless expression (magenta) detected by Bendless-specific antibody upon whole LG- LG-specific *e33cGal4*-mediated Bendless knockdown (C-D) or over-expression (E-F) as compared to the wildtype control (A-B). Nuclei are stained with DAPI (Blue). Scale Bar: 50µm (A-F).

**Figure S3: Hematopoietic progenitor- specific Bendless depletion or abrogation of K63 ubiquitination results in loss of progenitor/prohemocyte population**

Hematopoietic progenitor population marked by tepIV-GFP (Green) upon tepIV-Gal4 mediated depletion of Bendless (B, E) or expression of a non-functional ubiquitin mutant, Ubc.K63R (C, F) as compared to the wildtype control (A, D). Prohemocyte index per LG lobe represented graphically (G). Nuclei are stained with DAPI (Blue). GFP (Green) is driven by *tepIV-Gal4*. Scale Bar: 50µm (A-F). Statistical analysis was performed using Student's t-test with Welch's correction. Each data point denotes data from a single LG lobe. P-values denoted are as follows: **** (P<0.0001) and ** (P<0.01).

**Figure S4: *traf6* mutant allele shows defective LG blood cell homeostasis**

Posterior Signaling Center (PSC) cell numbers marked by Antennapedia (magenta), plasmatocyte differentiation marked by P1 (magenta), or crystal cell differentiation marked by Hindsight (Hnt, magenta) in the $traf6^{ex1}$ mutant allele (B, E, H) as compared to wild type control (A, D, G). Graphical representation of PSC cell numbers, plasmatocyte differentiation index or number of crystal cells (C, F, I) in the $traf6^{ex1}$ mutant as compared to wild type control. Nuclei are stained with DAPI (Blue). Scale Bar: 50µm (A-B, D-E, G-H). Statistical analysis was performed using Student's t-test with Welch's correction. Each data point denotes data from a single LG lobe. P-values denoted are as follows: **** (P<0.0001) and ** (P<0.01).

**Figure S5: Validation of K63 Ubiquitination abrogation and over-activation constructs**

Ub-K63 (magenta) expression upon *tepIVGal4* mediated expression of *UAS-UBC.K63R* (F-J) or *UAS-UBC.K63* (K-O) compared to wild type control (A-E). Nuclei are stained with DAPI (Blue). GFP (Green) is driven by *tepIV-Gal4.* Scale Bar: 50µm (A-B, F-G, K-L), 30µm (C-E, H-J, M-O).

**Figure S6: Dishevelled depletion in hematopoietic progenitors results in increased blood cell differentiation**

Plasmatocyte differentiation marked by P1 (magenta) or crystal cell differentiation marked by Hnt (magenta) upon *tepIVGal4* mediated Dishevelled knockdown using UAS-*dshRNAi* (B, E) in the tepIV positive hematopoietic progenitor population (green) as compared to wild type (A, D). Graphical representation of Plasmatocyte differentiation index or total number of crystal cells upon knockdown of Dishevelled in the progenitor population, compared to the wild type (C, F). Nuclei are stained with DAPI (Blue). GFP (Green) is driven by *tepIV-Gal4*. Scale Bar: 50µm (A-B, D-E). Statistical analysis was performed using Student's t-test with Welch's correction. Each data point denotes data from a single LG lobe. P-values denoted are as follows: **** ($P<0.0001$) and * ($P<0.1$).

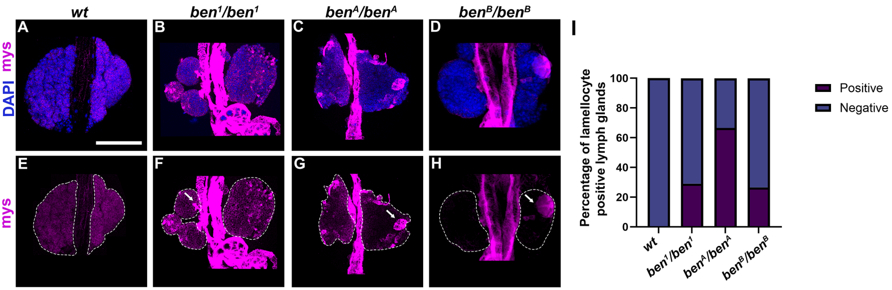

wt
ben1/ben1
benA/benA
benB/benB
A
B
C
D
E
F
G
H
I
DAPI mys
mys
Percentage of lamellocyte positive lymph glands
0
20
40
60
80
100
Positive
Negative

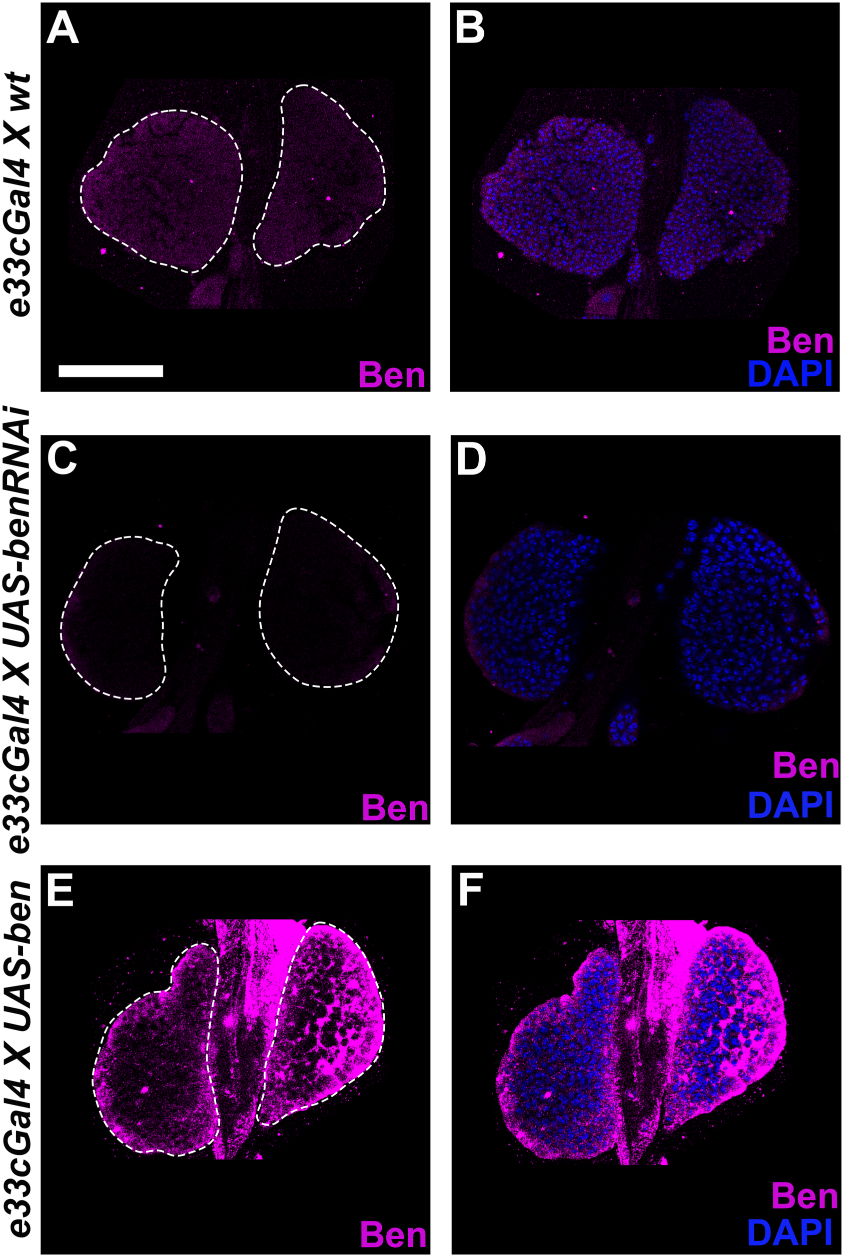

e33cGal4 X wt
A
Ben
B
Ben
DAPI
e33cGal4 X UAS-benRNAi
C
Ben
D
Ben
DAPI
e33cGal4 X UAS-ben
E
Ben
F
Ben
DAPI

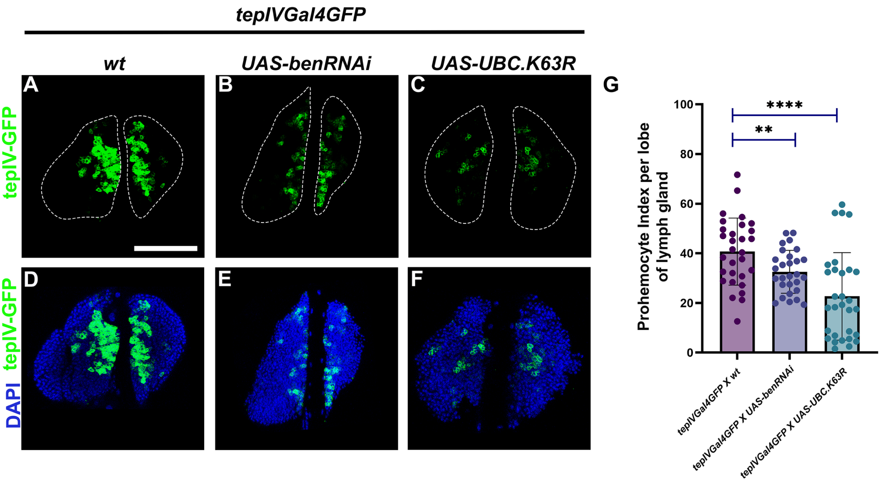

tepIVGal4GFP
wt
UAS-benRNAi
UAS-UBC.K63R
A
B
C
G
tepIV-GFP
D
E
F
tepIV-GFP
DAPI
Prohemocyte Index per lobe of lymph gland
100
80
60
40
20
0
****
**
tepIVGal4GFP X wt
tepIVGal4GFP X UAS-benRNAi
tepIVGal4GFP X UAS-UBC.K63R

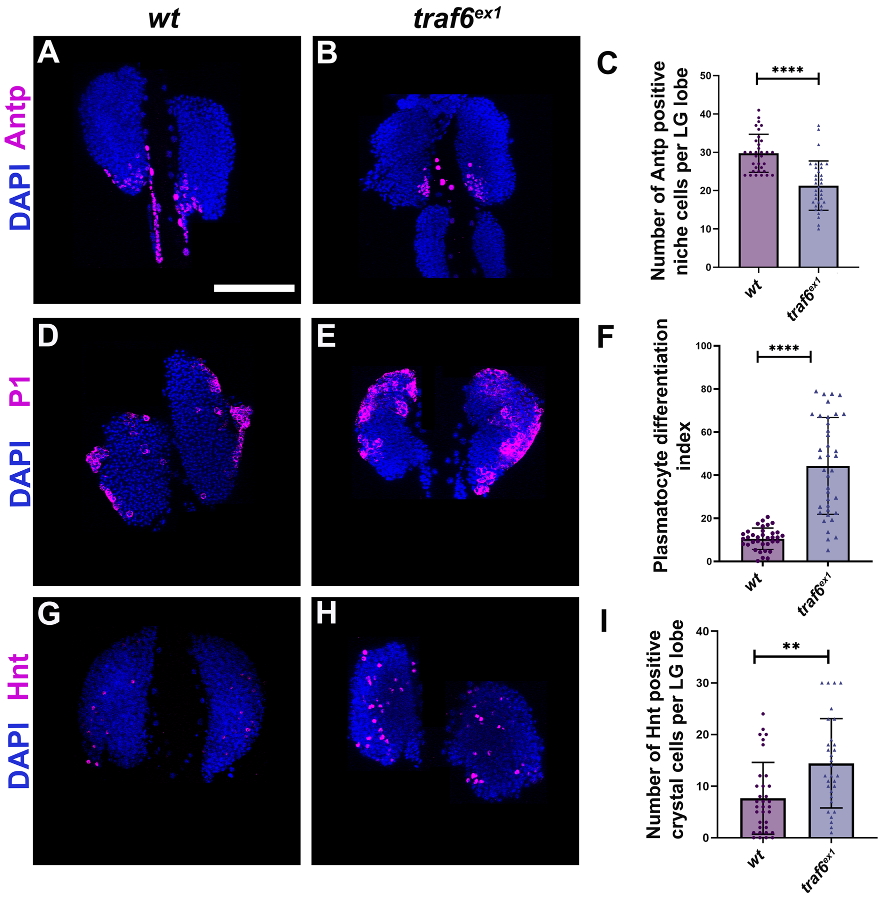
wt
traf6ex1
A
B
C
DAPI Antp
Number of Antp positive niche cells per LG lobe
****
D
E
F
DAPI P1
Plasmatocyte differentiation index
****
G
H
I
DAPI Hnt
Number of Hnt positive crystal cells per LG lobe
**

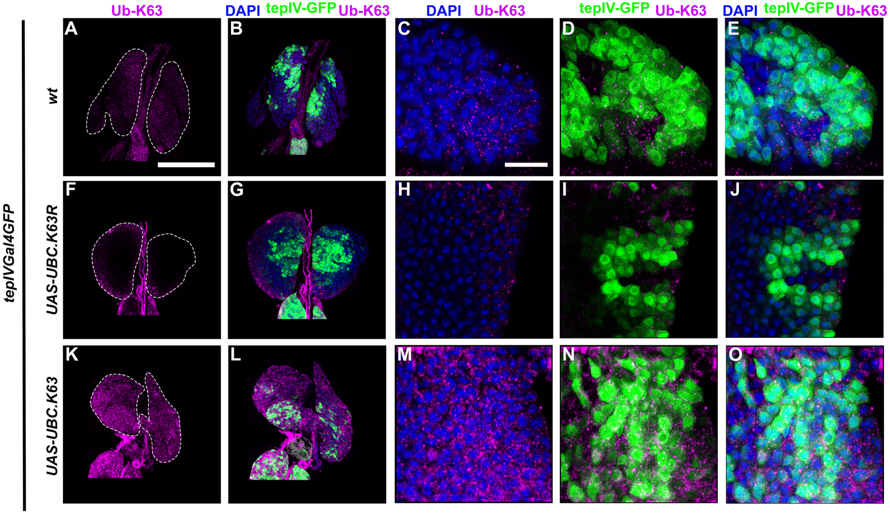
Ub-K63
DAPI tepIV-GFP Ub-K63
DAPI Ub-K63
tepIV-GFP Ub-K63
DAPI tepIV-GFP Ub-K63
A
B
C
D
E
F
G
H
I
J
K
L
M
N
O
wt
UAS-UBC.K63R
UAS-UBC.K63
tepIVGal4GFP

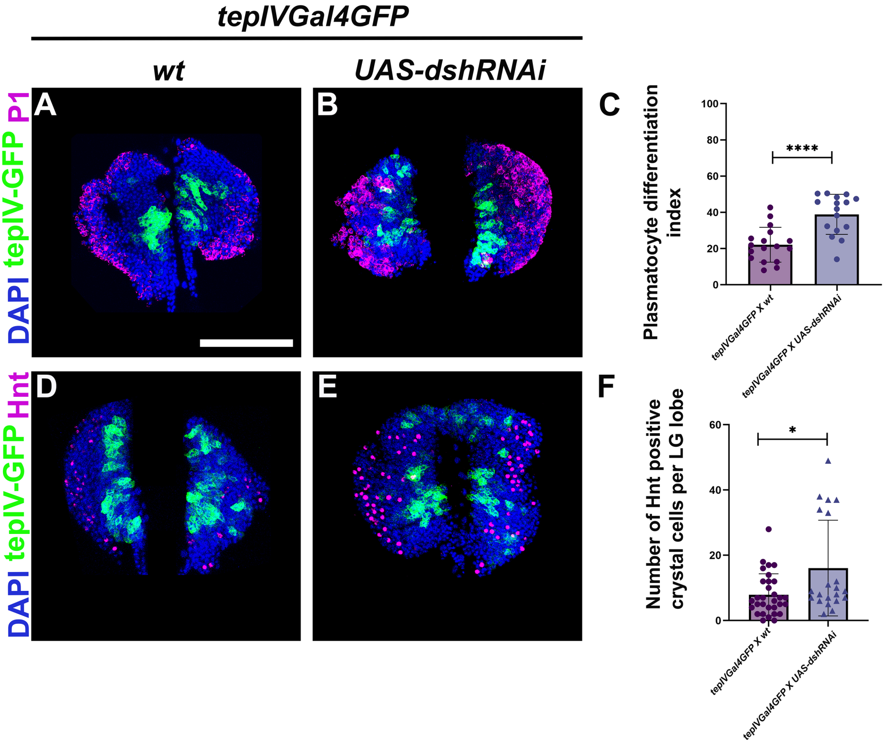

teplVGal4GFP
wt
UAS-dshRNAi
DAPI teplV-GFP P1
A
B
C
Plasmatocyte differentiation index
100
80
60
40
20
0
****
teplVGal4GFP X wt
teplVGal4GFP X UAS-dshRNAi
DAPI teplV-GFP Hnt
D
E
F
Number of Hnt positive crystal cells per LG lobe
60
40
20
0
*
teplVGal4GFP X wt
teplVGal4GFP X UAS-dshRNAi
